# Analysis *in vivo* using a new method, ARGO (Analysis of Red Green Offset), reveals complexity and cell-type specificity in presynaptic turnover of synaptic vesicle protein Synaptogyrin/SNG-1

**DOI:** 10.1101/2024.11.26.625560

**Authors:** Nikita Shiliaev, Sophie Baumberger, Claire E. Richardson

## Abstract

In long-lived cells such as neurons, proteostasis involves the regulated degradation and replacement of proteins to ensure their quality and appropriate abundance. Synaptic vesicle (SV) protein turnover in neurons is important for controlling the SV pool size to maintain appropriate levels of neurotransmission; yet, it is incompletely understood, partly due to limited tools for quantifying protein turnover in vivo. We present ARGO (Analysis of Red-Green Offset), a fully genetically encoded, ratiometric fluorescence imaging method that visualizes and quantifies protein turnover with subcellular resolution in vivo. ARGO is inexpensive, modular, and scalable for use in genetically tractable experimental organisms. Using ARGO, we examine the turnover of Synaptogyrin/SNG-1, an evolutionarily conserved, integral SV protein, in *C. elegans* neurons. We show that the SNG-1 turnover rate is consistent across presynapses within a single neuron but varies between neuron classes. Notably, we find SNG-1 and can exist in two distinct, non-intermixing populations within each presynapse. Further, we present an initial mutant analysis of *uba-1*, the sole E1 ubiquitin ligase in *C. elegans*, showing that we can detect slowed SNG-1 turnover even though steady-state SNG-1 abundance is not increased compared to wild-type. These results provide new hints for the regulation of SV pool size.

**Significance Statement:**

- In long-lived cells, a protein’s rates of synthesis and degradation together determine its abundance, yet regulation of protein turnover is largely unknown due to the lack of simple methods for in vivo quantification.
- Using *C. elegans*, the authors develop ARGO, a genetically encoded, microscopy approach that quantifies a protein’s turnover. Results suggest that synaptic vesicle protein Synaptogyrin/SNG-1 can partition into two presynaptic pools with distinct half-lives.
- This provides a powerful tool to study protein turnover, reveals an unexpected complexity in SNG-1 turnover, and lays the groundwork for future investigations of synaptic vesicle protein compartmentalization, sorting, and degradation.

## Introduction

For neurons and other long-lived cells, maintenance of cellular function requires maintenance of proteostasis. Part of proteostasis is the continuous turnover and replacement of most proteins, which is proposed to generate multiple benefits; these include promoting stability in protein levels despite stochastic variations and errors in gene expression, removing damaged proteins, and minimizing accumulation of protein damage, and regulating protein abundance (Reddien, 2024). The importance of this continuous turnover is highlighted by the consequence of its failure: declining proteostasis, including the accumulation of misfolded and damaged proteins, is a hallmark of aging and a central driver of neurodegenerative disease (Hipp *et al*., 2019; López-Otín *et al*., 2023). Neurons’ complex morphology presents a spatial challenge for turnover of synaptic proteins, as most protein degradation machinery resides in the neuron cell body (Winckler *et al*., 2018). However, homeostatic protein degradation remains incompletely understood for most proteins in any cell type. An obstacle to understanding this is the lack of easy and robust methods to interrogate protein half-life *in vivo* with high spatial and temporal resolution.

Several methods can be used to quantify homeostatic protein turnover. Pulse-chase labeling with heavy-isotope-containing amino acids followed by bulk proteomics to measure reduction in labeled protein over time has the strength of simultaneously measuring the half-life of thousands of proteins (Zhou, 2004). This method is laborious, though, especially *in vivo,* and the ability to measure protein half-life with cellular or subcellular resolution is limited since all of an animal’s cells are labeled. Fluorescence microscopy-based approaches enable assessment of an individual protein-of-interest’s turnover with high resolution. This has been achieved using photoconvertible fluorescent proteins, which requires calibration of the photoconversion to minimize phototoxicity in the pulse (Zhang *et al*., 2007). Another strategy is to label the protein-of-interest with a self-labeling tag, most commonly SNAP-tag or HaloTag, provide a “pulse” of fluorescent ligand, then image turnover with fluorescence microscopy during the “chase” (Bojkowska *et al*., 2011). Such experiments contributed evidence that, in cultured hippocampal neurons, SV proteins function in SV cycling for less than half of their total half-life, and, once decommissioned from SV cycling, they continue to reside at presynapse (Truckenbrodt *et al*., 2018).

Considering those results, one might speculate that the management of SV protein turnover as a whole, including the sorting out of the functional pool, is especially stringent Recently, this method was used to visualize and quantify the homeostatic degradation of post-synaptic protein PSD95 in the mouse brain using PSD95-HaloTag (Bulovaite *et al*., 2022). This was the first time homeostatic protein degradation of any neuronal protein has been quantified with notable spatial resolution *in vivo*, and the results indicated that PSD95 turnover depends on animal age, region of the brain, and subcellular region within individual neurons (Bulovaite *et al*., 2022). Such variation is consistent with the notion that synaptic protein turnover is highly regulated and may play in important role in specifying differences in synaptic function across different neuron types.

Importantly, comparing protein half-life between multiple separate studies that used the heavy-isotope-pulse-plus-proteomics approach shows that cultured neurons have dramatically shorter protein half-lives overall than neurons *in vivo* (Price *et al*., 2010; Cohen *et al*., 2013; Visscher *et al*., 2016; Dörrbaum *et al*., 2018; Fornasiero *et al*., 2018; Heo *et al*., 2018; Mathieson *et al*., 2018; Kluever *et al*., 2022). This indicates that there is a difference in the regulation of protein degradation in an adult animal’s functioning nervous system versus in immature, cultured neurons. This could be due to a variety of factors, including neuron age, neuron morphology, or cell-extrinsic environment; nevertheless, it highlights the value of studying protein turnover *in vivo*.

Here, we developed a new method to visualize the turnover of a protein of interest with high spatial and temporal resolution *in vivo* using fluorescence microscopy. We call this method ARGO (Analysis of Red Green Offset), drawing an analogy to the Theseus Ship Paradox, which ponders how identity persists even as components are gradually replaced over time. This method could, in theory, be applied to any protein-of-interest for which addition of a fluorescent tag does not impact localization or function.

We used ARGO to begin to directly analyze the turnover of SV proteins in adult neurons *in vivo.* SV proteins of are particular interest because the abundance, composition, and functionality of SV pools at each presynapse contributes to determining the amount of neurotransmission. Furthermore, measuring their turnover provides an opportunity to test whether SV pools are maintained as spatially or functionally distinct compartments within individual neurons, as non-uniform intracellular turnover would imply such distinctions. We chose Synaptogyrin/SNG-1 as our first protein of interest because it is an evolutionarily conserved transmembrane SV protein, it is present on most SVs, and it regulates SV cycling (Abraham *et al*., 2011; Raja *et al*., 2019). These features make SNG-1 an informative starting point for establishing ARGO and for probing principles of SV protein turnover. As it is an integral membrane protein, it is expected to be degraded via endosomal sorting to lysosomal compartments.

Indeed, prior research mainly using assessment of steady-state SV protein localization combined with genetic analyses has led to the model that SV proteins are sorted into late endosomes and/or autophagosomes at the synapse, then transported to the cell body for degradation within lysosomal compartments (Uytterhoeven et al. 2011; Fernandes et al. 2014; Sheehan et al. 2016; Okerlund et al. 2017). Interestingly, though, an imaging strategy to assess steady-state SV protein degradation during brain development in *D. melanogaster* found evidence that at least some SV protein is degraded in an axonal degradative “hub” (Jin *et al*., 2018). Another aspect under investigation is how the rate of SV protein degradation is regulated, with evidence pointing to both surveillance and/or sorting for degradation as rate-limiting steps (Fernandes *et al*., 2014; Sheehan *et al*., 2016; Okerlund *et al*., 2017; Sheehan and Waites, 2019; Hoffmann-Conaway *et al*., 2020; Birdsall *et al*., 2022). A more direct observation of SV protein turnover will potentiate a deeper understanding of this process.

## Results

### Strategy and validation of SNG-1::ARGO

To enable the analysis of protein turnover with subcellular resolution *in vivo*, we developed the ARGO (Analysis of Red Green Offset) method (Figure 1A). The ARGO construct cell-specifically, co-translationally labels the protein-of-interest with both RFP and GFP in a tandem tag. Cell specificity is achieved using FLP/FRT, wherein the Flippase is expressed from cell-specific promoters. The steady-state GFP/RFP ratio of an ARGO-tagged protein reflects the mechanism of degradation. As GFP fluorescence is quenched in acidic environments, ARGO-tagged proteins that have been sorted for degradation within lysosomal compartments are detected as endosomes with stronger RFP than GFP fluorescence (Figure 1B). Alternatively, the absence of detectable lysosomal compartments suggests that the protein-of-interest is degraded by the proteasome, or not at all. Next, the ARGO “pulse” removes *gfp* from the *argo* gene cassette such that all newly produced molecules of the protein-of-interest are labeled with only RFP. The neuron is then periodically imaged with fluorescence confocal microscopy, and the GFP/RFP ratio is quantified for each fluorescent punctum throughout the cell to determine the proportion of “old” versus “total” protein-of-interest both spatially and temporally.

**Figure 1.**
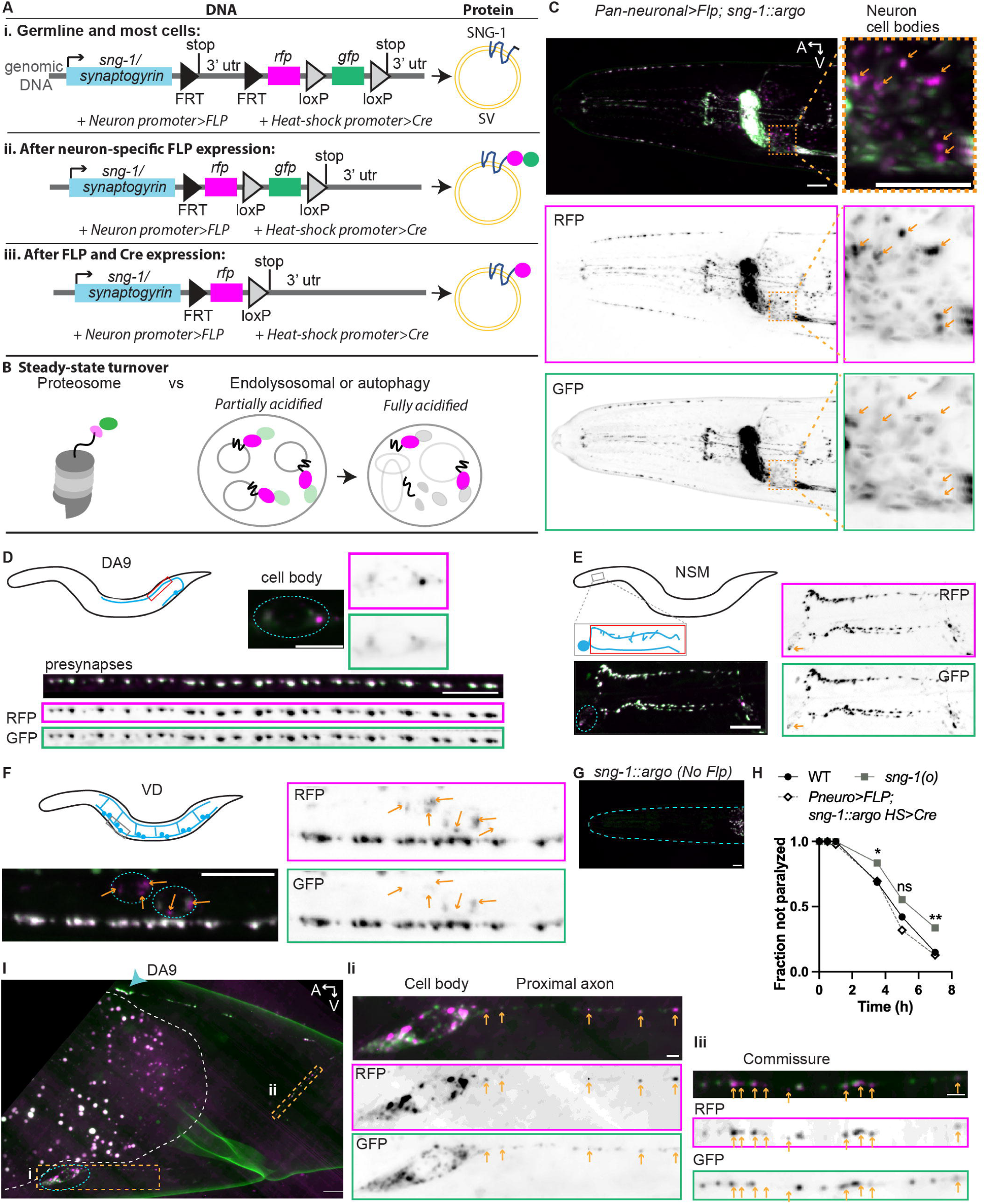
Steady-state SNG-1::ARGO fluorescence indicates that SNG-1 is degraded in lysosomal compartments in the cell body. (A-B) ARGO method overview. (Ai) The argo gene cassette is inserted into the gene-of-interest, ideally at the endogenous genomic locus. The strain also contains transgenes encoding neuron-specific FLP (Flippase) and a mechanism for inducible Cre expression. In this case, we tagged sng-1 at the genomic locus using CRISPR/Cas9 and used a heat-shock promoter to induce Cre expression. (right) The protein-of-interest, in this case SNG-1, will be expressed with a short additional amino acid tag, which is encoded by the first FRT (Figure 1 continued) sequence, in every cell. (Aii) Neuron-specific promoter drives Flp expression shortly after neuron birth. These cells “turn on” the ARGO tag. (Aiii) (left) A pulse of Cre expression removes gfp from the argo gene cassette. This “activates” the ARGO. (right) All newly synthesized SNG-1 will be tagged with RFP. Therefore, RFP fluorescence reports total SNG-1 protein, whereas GFP reports “old” SNG-1. (B) Steady-state imaging provides clues as to how, and sometimes where, a protein-of-interest is degraded. For ARGO-tagged proteins that are degraded in lysosomal compartments, protein that have been sorted for degradation will show stronger RFP vs GFP fluorescence, as GFP is quenched by the acidic environment of the endolysosomal and/or autolysosomal lumen. (C) Representative image of the head of an animal with the sng-1::argo allele in which the ARGO tag is flipped-on by a pan-neuronally expressed FLP, showing co-localized RFP and GFP fluorescence at presynapses throughout the nervous system. Zoomed-in views on the right shows a region where neuron cell bodies reside. Arrows point to endosomes with relatively brighter RFP vs GFP intensity. Scale = 10 18lm (D-F) SNG-1::ARGO Flipped-on in individual neurons. Cartoons show neuron morphology (blue) within the animal (black). Red box shows where presynapses are located (D-E), gray boxes indicate the region imaged (E-F), and teal dashed ovals encircle neuron cell bodies (D-F). The VD neurons (F) make a string of en passant synapses along the ventral nerve cord. (G) In the absence of FLP expression, there is no detectable SNG-1::ARGO fluorescence. Teal dashed curve outlines the nose of the animal. (H) Sensitivity to 0.5 mM aldicarb in Day 1 adults (A1) (*P<0.05, **P<0.01, Chi-squared test with Bonferroni correction, n = 90 animals per genotype summed across three separate experiments). (I) SNG-1::ARGO in the DA9 neuron imaged with Super Resolution spinning disk confocal microscopy with deconvolution. Area inside dashed white line contains autofluorescence from the intestine. Teal dashed oval encircles the neuron cell body; teal arrowhead points to the most proximal presynapse. Note that transport packets containing SNG-1::ARGO are visible throughout the axon between the cell body and the proximal synapse. Scale= 5 um. (Ii-ii) Transport packets with brighter RFP vs GFP fluorescence are indicated with orange arrows. Scale = 1 um.

To study SV protein turnover, the *argo* cassette was inserted into the *sng-1* genomic locus such that the RFP::GFP tandem tag will be attached to the C-terminus of SNG-1, which resides on the cytoplasmic side of SVs (Figure 1A). Expression of Flippase from a pan-neuronal reporter shows SNG-1::ARGO dually labeled with RFP and GFP fluorescence throughout the nervous system (Figure 1C). Driving Flippase expression from neuron-specific promoters, we turned on SNG-1::ARGO expression in three individual neuron classes that represent a variety of morphologies and functions: the cholinergic motor neuron DA9, the two bilaterally symmetric serotonergic NSM neurons, and the GABAergic motor neurons VD/DD (Figures 1D-F). Each of these neuron types elaborates multiple *en passant* synapses that can be visualized as individual puncta using fluorescence confocal microscopy (White *et al*., 1986). The use of neuron-specific Flippase transgenes enables visualization of these individual presynapses, to which SNG-1::ARGO localizes as expected, with co-localized fluorescence from RFP and GFP (Figures 1D-F)(Schwartz and Jorgensen, 2016; Muñoz-Jiménez *et al*., 2017). In the absence of Flippase, no fluorescence is observed from the *sng-1::argo* allele (Figure 1G).

For the ARGO method to provide meaningful information about protein turnover, it must not impact the function of the protein-of-interest. This is because each of the facets of a protein’s function, including folding, localization, and activity, are factors that could be inputs to regulate its turnover. For SNG-1::ARGO, proper localization to the presynapses is the first indicator that this allele is usable. Next, we assessed *sng-1* function. Synaptogyrin/SNG-1 regulates synaptic transmission from worms to mammals (Abraham *et al*., 2011; Raja *et al*., 2019). In *C. elegans*, loss of *sng-1* function causes resistance to the acetylcholinesterase inhibitor aldicarb, suggesting that the *sng-1(o)* mutant has reduced SV release compared to wild-type (Figure 1H)(Mahoney *et al*., 2006; Abraham *et al*., 2011). By contrast, pan-neuronally expressed SNG-1::ARGO has no discernable effect on aldicarb resistance (Figure 1H), suggesting that the SNG-1::ARGO protein is functional. Together, these results suggest that SNG-1::ARGO is likely a valid tool for investigating SNG-1 biology.

To generate the pulse that activates ARGO, we excised *gfp* from the *ARGO* gene cassette using a heat-shock promoter to drive production of Cre recombinase upon a brief heat shock. This is the quickest and most robust method to induce gene expression in *C. elegans*. Indeed, by assessing fluorescence in non-heat-shocked animals and in animals several days post-heat-shock, we observed that this method efficiently activated SNG-1::ARGO in each of the three neuron types we examined, with little ectopic excision of *gfp* in the absence of heat shock (Table 1, Figure S1A). Though it is practical, a drawback to this approach is that heat shock causes increased production of HSF-1-regulated genes including chaperones, which will impact proteostasis. To assess whether this is likely to have a substantial impact on SNG-1 turnover dynamics or mechanisms, we assessed whether *sng-1* mRNA or protein abundance are altered by the heat shock pulse. We reasoned that, if neither the rate of SNG-1 production nor the steady-state abundance of SNG-1 is impacted by the heat shock, then it is unlikely that the degradation is altered. We observe no significant difference in *sng-1* mRNA levels either 5 hours or 1 day after heat-shock (Figure S1B). To assess steady-state SNG-1 protein abundance, we quantified average presynaptic SNG-1::ARGO RFP intensity and number of synapses in the DA9 neuron. There was no significant difference 4 hours, 1 day, or 2 days after heat shock compared to control (Figure S1C-D). We detect an increase in the presynaptic SNG-1::ARGO GFP/RFP ratio immediately following heat-shock that is not detectable by 4 hours or 1 day post-heat shock (Figure S1E). That timing is expected to be too fast to reflect a change in *sng-1* gene expression; it is therefore likely to reflect a change in localization, such as decreased SV cycling or trafficking. Still, the difference is resolved by 4 hours post heat-shock and that the first post-heat shock imaging timepoint used in turnover experiments is 1 day. Together, these data indicate that SNG-1::ARGO turnover is unlikely to be substantially impacted by the heat shock pulse. Concerns about a potential subtle impact are mitigated by the fact that all turnover comparisons were made between cohorts that received the same heat shock.

**Table 1.** Specificity and Efficacy of SNG-1::ARGO Activation by Heat-shock-Promoter driving *Cre* recombinase.

| Specificity |  |  |
| --- | --- | --- |
| Neuron | Age scored (d of adulthood) | GFP not recombined without heat shock (n) |
| DA9 | >= 4 | 95% (146) |
| NSM | >=6 | 100% (43) |
| VD2 | >=5 | 99% (88) |
| Efficacy |  |  |
| Neuron | Age heat-shocked (d of adulthood) | GFP recombined after heat-shock (n) |
| DA9 | 0 | 97.3% (108) |
| DA9 | 2 | 94% (65) |
| NSM | 2 | 100% (55) |
| VD2 | 2 | 97% (38) |

Parameters were assessed in strains carrying *sng-1(syb3140car2[argo])* for DA9 and NSM and *sng-1(syb3140[unstable argo])* for VD2. All strains carried an individual neuron-specific flippase (one per strain), and an integrated transgene composed of heat-shock promoter driving *Cre* recombinase (*heSi160*). Specificity means how often the *GFP* was not excised in the absence of heat-shock, and Efficacy means how often the *GFP* was excised upon heat-shock of 1 h at 34 °C.

Experimentally, the ARGO pulse is a 1-hour heat shock at 34 °C, but the pulse in terms of generating a Time = 0 for turnover experiments requires production of Cre, Cre-mediated removal of the *gfp* gene, and degradation of the mRNA molecules encoding *sng-1::rfp::gfp.* We assessed the effective timing of the pulse by imaging presynapses 4 hours post-heat shock (Figure S1F-L). Whereas we detect no change in SNG-1::ARGO GFP/RFP in a heat-shocked strain lacking the *HS>Cre* transgene, there was a significant decrease in the GFP/RFP ratio in five of the six neuron-by-animal-age conditions tested (Figure S1F-L). In addition, the abundance of Cre-recombined *sng-1::argo* alleles in the genome peaked by 4 hours post-heat shock (Figure S1M). Therefore, the ARGO pulse effectively takes on the order of several hours, and the ARGO method systematically over-estimates protein half-life by that much. For comparison, the mean half-life of the *C. elegans* proteome is estimated to be on the order of a day or two (Depuydt *et al*., 2016; Visscher *et al*., 2016).

### SNG-1 is likely sorted for degradation at the presynapse and degraded in the cell body

Assessing steady-state SNG-1::ARGO, we observed that there are fluorescent puncta within the neuron cell bodies that show relatively stronger RFP intensity compared to GFP intensity (Figure 1C-F). Super-resolution microscopy resolved two distinct populations of SNG-1::ARGO-labeled endosome in the neuron cell body: GFP-brighter endosomes, which are likely newly synthesized and trafficking endosomes, and RFP-brighter endosomes, which are likely SNG-1 within acidic lysosomal compartments (Figure 1I). Furthermore, transport packets of both the GFP-brighter and RFP-brighter flavors are present along the proximal axon and commissure (Figure 1I). These data suggest that SNG-1::ARGO is trafficked retrogradely from the presynapses to the cell body in acidified lysosomal compartments for degradation (Figure 1I).

We measured the steady-state SNG-1::ARGO GFP/RFP ratio at each presynapse per neuron in the DA9 neuron from Day 0 (L4 larval stage, A0) to Day 7 of adulthood (A7) (Figure 2A). For comparison, mean adult lifespan is ∼15 days (Gems and Riddle, 2000). Note that GFP has a higher quantum yield than RFP and is consistently brighter *in vivo,* so a ratio greater than 1 was expected (Rodriguez *et al*., 2017). Indeed, mean presynaptic SNG-1::ARGO GFP/RFP ratio at A0 was >2 (Figure 2A). Mean presynaptic SNG-1::ARGO GFP/RFP declined between A0 to A2, then remained fairly stable, with a trend for progressive slight decrease that was resolved at A7 (Figure 2A).

**Figure 2.**
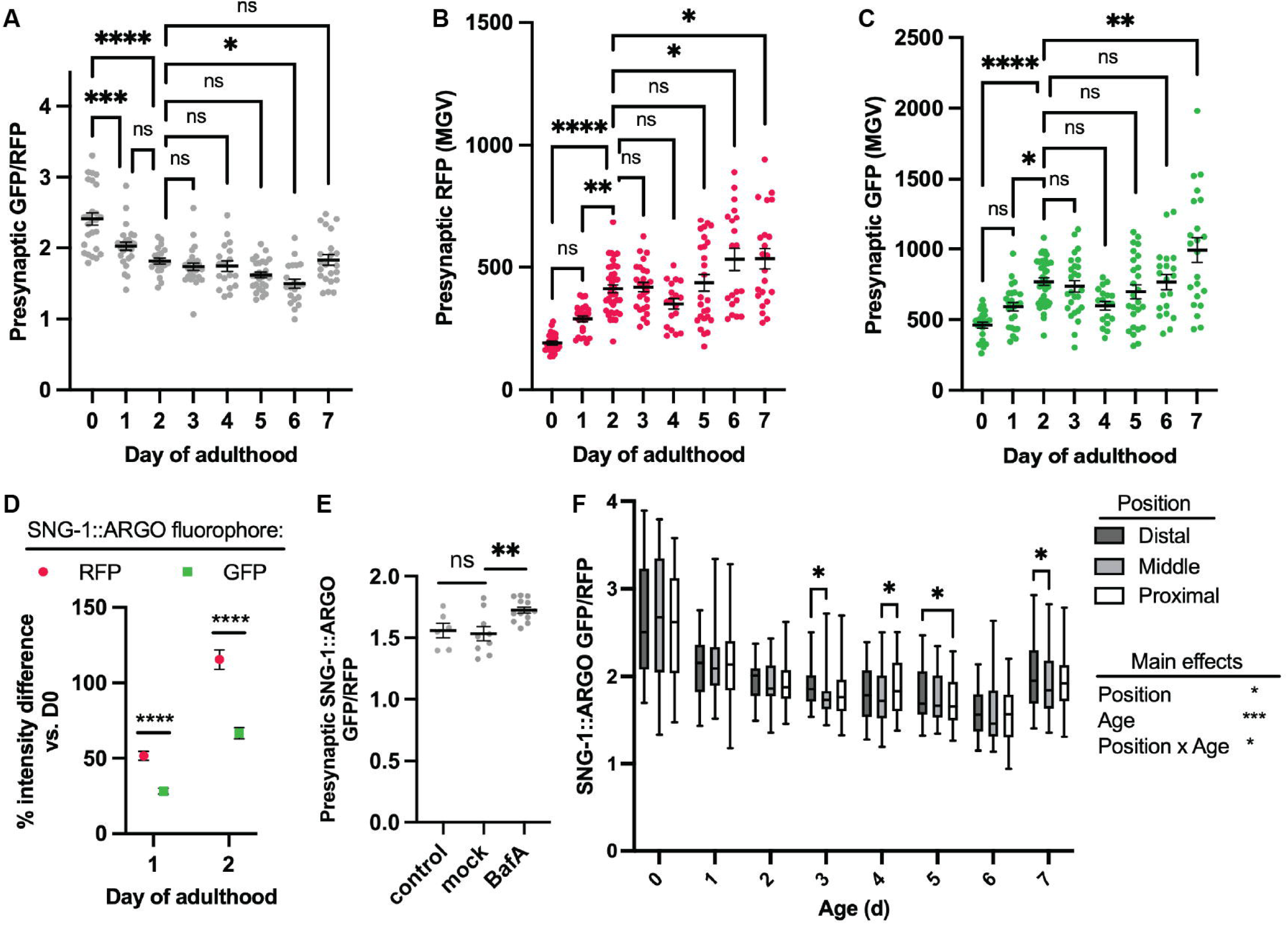
Presynaptic SNG-1::ARGO shows a decline in steady-state GFP/RFP fluorescence, indicative of accumulation within acidic compartments, during early adulthood in the DA9 neuron. (A-C) Average presynaptic RFP/GFP ratio (A), and the underlying intensity data for RFP (B) and GFP (C) from SNG-1::ARGO. Data points are each the mean of all presynapses from a single neuron. Here and elsewhere unless otherwise noted, mean± SEM are plotted. (D) Percent change in average presynaptic RFP and GFP intensity from Day 0 (AO) through Day 2 (A2) of adulthood, calculated from the data in B-C. (E) Presynaptic SNG-1:ARGO GFP/RFP in A3 animals injected with Bafilomycin A (BafA), vehicle (“mock”), or uninjected (“control”), quantified and displayed as in A. (F) SNG-1::ARGO presynaptic GFP/RFP by presynapse position along the axon. ns: not significant, *P<0.05, **P<0.01, ***P<0.001, ****P<0.0001, one-way ANOVA (A-C,E) or two-way ANOVA (D,F) with Tukey post-test.

To better understand the change in ratio between A0 to A2, we examined the RFP and GFP intensities underlying the ratiometric data (Figure 2B-C). *C. elegans* body size increases between A0 and A2, and the DA9 synapses likewise increase in both number and size based on the RFP intensity (Figure S1D, 2B, 2D). The average steady-state (no pulse) SNG-1::ARGO GFP fluorescence intensity per synapse likewise increases, but by only half as much, underlying the decrease in the presynaptic SNG-1::ARGO GFP/RFP ratio (Figure 2C-D).

We hypothesized that A) SNG-1::ARGO is sorted into acidified compartments, likely late endosomes, at the presynapse, which partially quenches the GFP fluorescence, and B) the steady-state proportion of presynaptic SNG-1::ARGO that resides within acidified compartments increases in early adulthood. To test this hypothesis, we injected mid-adult animals with Bafilomycin A, which inhibits the V-ATPase that acidifies lysosomes and late endosomes. Indeed, animals injected with Bafilomycin A showed an increase in the presynaptic SNG-1::ARGO GFP/RFP ratio 3 hours post-injection compared to control and vehicle-injected animals (Figure 2E). Furthermore, we have shown that injecting animals with concanamycin A, which also blocks the V-ATPase, increased the SNG-1::ARGO GFP/RFP ratio at both the presynapse and the neuron soma within hours (Zhong and Richardson, 2025). Consistent with this interpretation, late endosomes and autophagosomes, which both acquire acidified lumens, localize to presynapses (Ceccarelli *et al*., 1973; LaVail and LaVail, 1975; Grill *et al*., 2007; Binotti *et al*., 2014; Soukup *et al*., 2016; Stavoe *et al*., 2016; Neisch *et al*., 2017).

Taken together, these data suggest that SNG-1::ARGO is sorted for degradation into acidified compartments at the presynapse and then trafficked to the cell body to complete degradation. This aligns with the prevailing model for SV protein turnover, which was mainly based on more indirect evidence including genetic analysis of steady-state abundance and the presence of MVBs and autophagosomes at presynapses. Of note, SNG-1 within partially acidified compartments is not necessarily destined for degradation; this notion is considered further below.

Next, we assessed whether presynaptic SNG-1::ARGO GFP/RFP ratio is dependent on presynapse position. We binned the presynapses as “proximal,” “middle,” or “distal” along the axon and compared the steady-state GFP/RFP for each day (Figure 2F). There was no consistent relationship between synapse position and ratio (Figure 2F).

### Quantification of presynaptic SNG-1::ARGO turnover in the DA9 neuron

Using the DA9 neuron, we pulsed SNG-1::ARGO at A2 and then imaged unique cohorts of animals daily for five days afterward, plus a cohort immediately before and after the activation (Figure 3A-B, S1H). SNG-1::ARGO GFP/RFP ratios were normalized to the mean ratio from age-matched control (non-pulsed) cohorts imaged in parallel. From A2, presynapse size (based on SNG-1::RFP intensity) and presynapse number are fairly stable (Figure 2B, S1A,C,D), meaning that the SNG-1::RFP level meaning that the declining SNG-1::ARGO GFP/RFP each day after the pulse is predominantly due to homeostatic SNG-1 turnover. Fitting a single-phase exponential decay function to the data calculated an apparent presynaptic half-life of 3.0 days (95% C.I. 2.7 - 3.4 days). We observed no impact of presynapse position relative to the neuron cell body on the half-life (Figure 3C). To assess whether some other feature of individual presynapses might regulate turnover rate within a neuron, we quantified the intraneuronal coefficient of variation (CV) in presynaptic SNG-1::ARGO GFP/RFP for each timepoint in the pulse-chase (Figure 3D). If the rate of SNG-1::ARGO turnover varies by presynapse, we expect to observe an increase in CV during the chase, especially at timepoints close to the pan-synaptic half-life, compared to T=0. However, no change in CV was detected.

**Figure 3.**
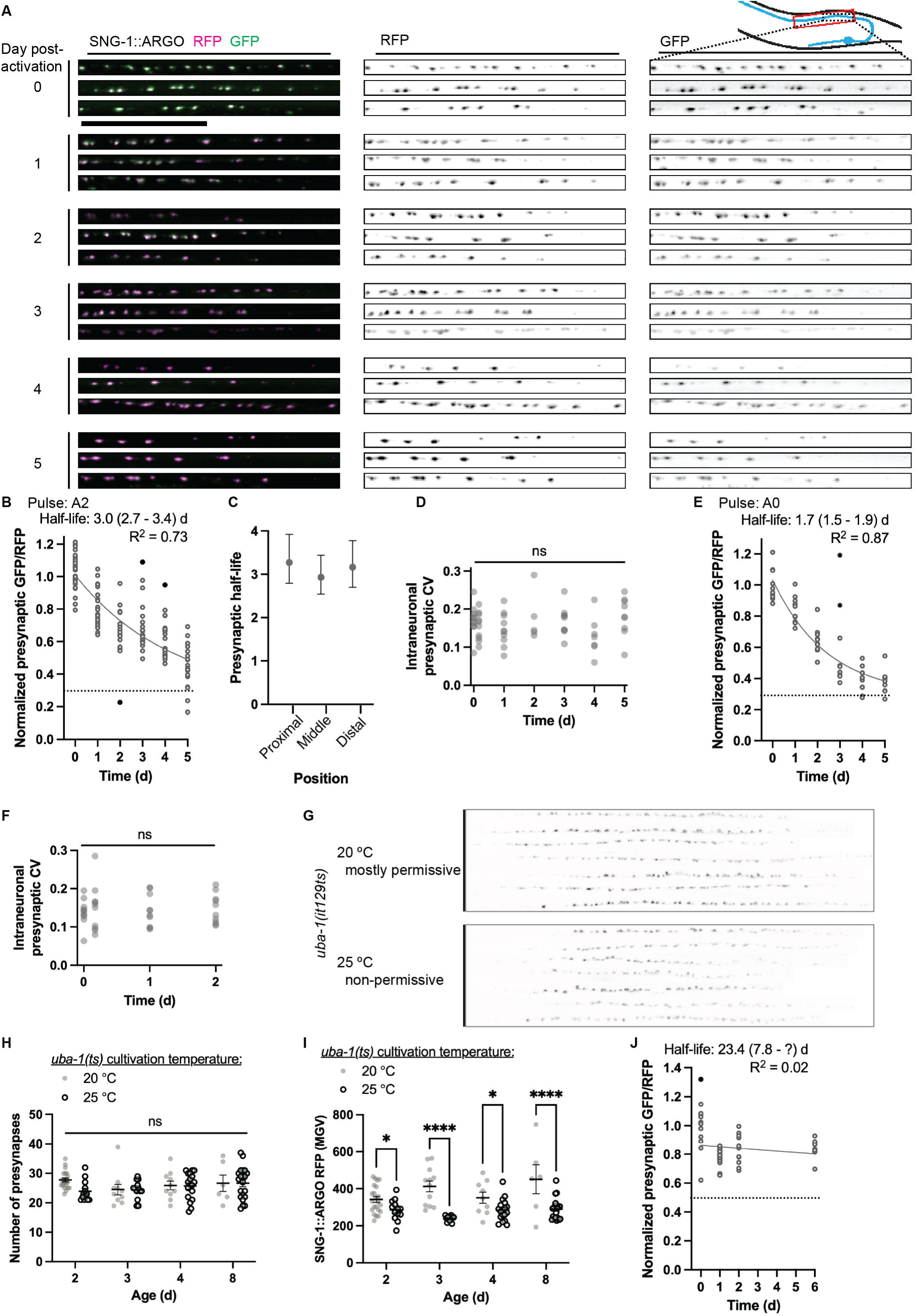
Quantification of SNG-1::ARGO turnover in the DA9 neuron. (A-B) Example images (A) and quantification (B) of SNG-1::ARGO in DA9 presynapses from turnover experiment with the pulse at A2. (A) Proximal synapses from three animals per time-point (different cohorts of animals are imaged at each timepoint). Scale = 30 µm. (B) One-phase exponential decay function fit to SNG-1::ARGO turnover data with the pulse at A2; each data point shows the average presynaptic SNG-1::ARGO GFP/RFP ratio from all the presynaptic puncta in a single neuron. Dashed black line shows the experimentally calculated background, which comes from autofluorescence. Outlier data points that were excluded from the curve calculation are shown in black. (C) SNG-1::ARGO presynaptic half-life by proximal-distal position relative to the neuron cell body (mean± 95% C.I.). (D) The average coefficient of variation (CV) for all the presynaptic puncta within each neuron does not change at any chase timepoint compared to the pulse, suggestive that there is effectively a uniform SNG-1::ARGO turnover rate across all presynapses within one neuron. (E-F) From a turnover experiment with the pulse at AO, one-phase exponential decay function (E) and intraneuronal variance between synapses (F). (G-I) The *uba-1(it129ts)* mutant at the non-permissive temperature, 25 C, shows no obvious defect in presynapse localization (G) or number (H) but reduced abundance of SNG-1::ARGO at each presynapse (I). Two-way ANOVA with Tukey post-test. (J-K) A turnover experiment started at adult Day 2 indicates that the SNG-1::ARGO presynaptic turnover is slowed in the *uba-1(it129ts)* mutant (J).

We performed a pulse-chase experiment with the DA9 SNG-1::ARGO starting at A0. From A0 to A2, the DA9 presynapses grow by over 100%, and additional presynapses are added, scaling with increasing animal size (Figure 2B-D, S1D). The apparent presynaptic SNG-1::ARGO half-life – 1.7 days (95% C.I. 1.5-1.9 days) – therefore measures a combination of both the removal of old molecules of SNG-1 from synapses and the increasing total abundance of SNG-1 (Figure 3E). The shorter apparent half-life with the pulse at A0 compared to A2 is likely due to this scaling as opposed to a change in the rate of SNG-1 degradation. Indeed, fitting one-phase exponential decay curves to both the A2 and A0 datasets using the GFP intensity rather than the GFP/RFP ratio, we observe no difference in apparent half-life between the two different pulse times (Figure S2A-B). With the pulse at A0, the calculated SNG-1 half-life is longer using mean GFP, which only quantifies degradation, compared to using the GFP/RFP ratio which quantifies degradation plus synapse growth. By contrast, with the pulse at A2, the half-life calculated using mean GFP is indistinguishable from that calculated using the GFP/RFP ratio, though the 95% C.I. is substantially larger. Indeed, using the GFP intensity instead of the GFP/RFP ratio to fit the exponential decay function results in substantially lower R^2^ values (>0.72 for ratiometric versus <0.29 for GFP only) and wider 95% C.I.s (<0.8 d for ratiometric versus >2.4 d for GFP only) for both experiments (Figure 3B-D, S2A-B). This demonstrates the utility of incorporating ratiometric quantification for turnover analysis.

Returning to the presynaptic scaling that occurs between A0 to A2, the SNG-1::ARGO pulse-chase experiment affords an opportunity to assess whether newly synthesized SNG-1::ARGO, which is tagged with only RFP, is allocated to the presynapses with any spatial pattern. Such a pattern could be generated by synapse position or activity (i.e. the amount of SV cycling), and it could be detected as an increase in the intraneuronal CV in presynaptic SNG-1::ARGO GFP/RFP ratio in the chase compared to the pulse. No such increase was detected, indicating that newly synthesized SNG-1 is evenly distributed throughout the neuron’s presynapses (Figure 3F). This is consistent with the model that SV deposition at both newly forming and fully formed presynapses strongly depends on trafficking parameters, and continuous trafficking intermixes SVs between synapses (Staras *et al*., 2010; Herzog *et al*., 2011; Wu *et al*., 2013; Maeder *et al*., 2014; Lipton *et al*., 2018).

### SNG-1::ARGO turnover is slowed in a *uba-1*/E1 Ubiquitin ligase loss-of-function mutant

As an initial mutant analysis using the ARGO method and a first step toward defining the mechanisms of SNG-1 turnover, we constructed a strain carrying the DA9 *sng-1::argo* alleles and a temperature-sensitive hypomorphic allele of *uba-1(it129ts),* which encodes the sole *C. elegans* E1 ubiquitin ligase (Kulkarni and Smith, 2008). Ubiquitination marks proteins for degradation via the proteasome, the endolysosomal system, and some types of autophagy (Foot *et al*., 2017; Yin *et al*., 2020). Animals cultivated at the non-permissive temperature, 25 °C, looked fairly wild-type in terms of presynaptic localization, organization, and number (Figure 3G-H), but they showed a reduction in presynaptic SNG-1::ARGO RFP intensity, indicating that *uba-1* is required to promote retention and/or delivery of SNG-1::ARGO to the presynapse (Figure 3I). In a turnover experiment with the pulse at A2, the *uba-1(it129ts)* mutant showed a presynaptic SNG-1::ARGO half-life of 23.4 days (95% C.I. 7.8-[upper limit could not be calculated])(Figure 3I, see also Figure S2C).

These data suggest that ubiquitination is likely required to promote SNG-1 degradation from the presynapse. One hypothesis is that SNG-1 gets ubiquitinated and then sorted for degradation via ESCRT; an alternate hypothesis is that SNG-1 turnover is regulated indirectly by ubiquitination of other protein(s) in the neuron. Because the *uba-1(it129ts)* mutant is hypomorphic rather than amorphic, it is possible that UBA-1 is essential for all presynaptic SNG-1 turnover. Of note, one might predict that a reduction in homeostatic protein degradation could be revealed with steady-state imaging as an increased abundance of the protein-of-interest; however, our results suggest that both presynaptic SNG-1 removal and steady-state abundance decrease in the *uba-1(it129ts)* mutant, highlighting the utility of quantifying turnover.

### Steady-state presynaptic SNG-1::ARGO varies with neuron identity

We assessed steady-state presynaptic SNG-1::ARGO GFP/RFP in the NSM and VD2 neurons. First, we compared the intraneuronal coefficient of variation across all presynaptic puncta of cohorts of DA9, NSM, and VD2 neurons at A2. If any neuron type showed a notably higher interneuronal CV, it could indicate that there is differential regulation in the sorting of SNG-1::ARGO into acidic compartments within individual presynapses. However, the NSM interneuronal CV was similar to that of DA9, and the CVs within VD2 neurons were low (Figure S2D).

We assessed steady-state SNG-1::ARGO across early adulthood and performed a turnover experiment with the pulse at A2. The SNG-1::ARGO steady-state and turnover characteristics generally were similar for NSM to those of the DA9 neuron (Figure S2D-J). The NSM neuron showed a stronger steady-state position effect, though, with a trend that the more proximal synapses had a lower SNG-1::ARGO GFP/RFP ratio compared to the more distal synapses (Figure S2H). Perhaps this is indicative of an increased density of SNG-1-containing lysosomal compartments closer to the cell body, which could arise due to SNG-1 retrograde trafficking in acidified compartments.

### Biphasic turnover of SNG-1::ARGO within each VD2 presynapse

In the VD2 neuron with SNG-1::ARGO activation at A2, a one phase exponential decay curve does not fit the data as well as it does for the DA9 and NSM neurons (VD2 R^2^ = 0.41, DA9 R^2^ = 0.73, NSM R^2^ = 0.63)(Figure 3B, 4A, S2I). A two-phase exponential decay curve fits the VD2 data better (R^2^ = 0.48), and it is apparent that the curve plateaus above the calculated background (Figure 4A). Indeed, according to the best-fit parameters for the two-phase curve, 39% of the SNG-1::ARGO molecules turn over with a half-life of about 0.9 days, and the other 61% has essentially an infinite half-life (Figure 4A). We considered whether this two-phase turnover could be due to non-specific or ineffective Cre-mediated recombination specifically for the VD2 neuron, i.e., either some VD2 neurons would fail to recombine one or both *gfp* alleles after the pulse and/or some VD2 neurons would recombine one or both alleles of *gfp* during steady-state imaging. To assess this idea, we evaluated the interneuronal CV for each neuron-by-age-by-treatment (Figure S3). If the VD2 neuron had inefficient and/or nonspecific removal of *gfp,* we would expect to observe a higher CV in the VD2 neuron during the pulse-chase experiment and/or steady-state imaging compared to the DA9 and NSM neurons. This trend was not observed (Figure S3).

**Figure 4.**
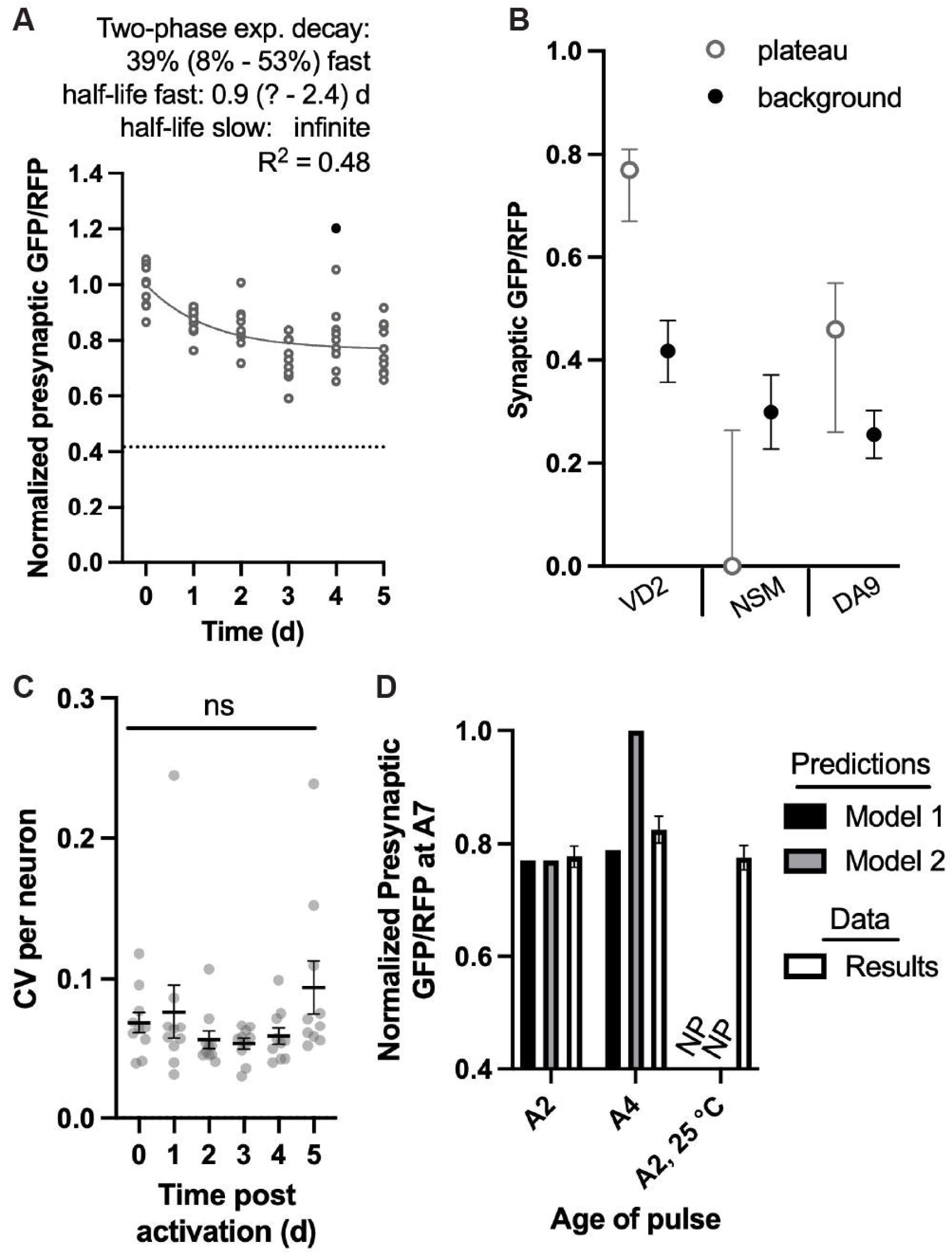
Biphasic SNG-1::ARGO turnover in the VD2 neuron. (A) SNG-1::ARGO turnover experiment in the VD2 neuron with the pulse at A2. Best-fit parameters and curve for a two-phase exponential decay function are shown, as this model was significantly favored over a one-phase exponential decay. (B) Comparison of the one-phase exponential decay function plateau (best-fit value and 95% C.I.) versus the background calculated from the images (mean± 95% C.I.) for each neuron identity with the pulse at A2. Data are from the experiments shown in 5A (VD2), S2I (NSM), and 3B (DA9). (C) lntraneuronal variance between individual presynaptic puncta during the VD2 turnover experiment. Each data point shows the CV for all the presynaptic puncta from one neuron. (D) Predicted presynaptic GFP/RFP values from the two models, qualitatively compared to results from a separate experiment in which animals with VD2 neuron SNG-1::ARGO were pulsed at different ages/conditions and imaged at A?. NP: no prediction.

As an alternate approach to assess whether the one-phase exponential decay curve can be ruled out as a good descriptor of VD2 SNG-1::ARGO turnover, we fit a one-phase exponential decay curve to the data with an unconstrained plateau. The best-fit plateau was significantly higher than the calculated background, and the 95% C.I.s from the two calculations did not overlap (Figure 4B). By the same analysis, the best-fit plateau and 95% C.I. for the NSM and DA9 neurons overlaps with that of the calculated backgrounds (Figure 4B).

We assessed whether the two-phase curve in the VD2 neuron arose from distinct turnover kinetics at spatially separate presynaptic puncta, i.e., some presynaptic puncta show turnover with a 0.9 day half-life while other presynaptic puncta within the same neuron show effectively no turnover. To do this, we calculated the CV between all the presynaptic SNG-1::ARGO GFP/RFP ratios within each neuron, across each timepoint of the turnover experiment (Figure 4C). We found no significant increase in the average CV at any day post-activation compared to T=0 (Figure 4C). These data support the model that each presynaptic punctum within an individual VD2 neuron exhibits the biphasic turnover.

We considered two models to explain the biphasic turnover in the VD2 presynapses. In Model 1, there are two chronically distinct populations of SNG-1::ARGO within each presynapse, one with a relatively fast turnover and the other with an unappreciable turnover. In Model 2, the mechanisms that promote SNG-1 turnover are shut off or become dysfunctional as the animal ages, at an age that is spanned by the turnover experiment. Model 1 predicts that the proportion of SNG-1 with an unappreciable turnover will be the same regardless of animal age at which the SNG-1::ARGO is activated (Figure 4D). By contrast, Model 2 predicts that the proportion of SNG-1 with an unappreciable turnover will be lower when SNG-1::ARGO is activated in younger animals and higher when it is activated in older animals (Figure 4D)(see Methods). To test these models, we performed an experiment in which we activated the VD2 SNG-1::ARGO at A2 (versus A4 Day 2 and Day 4 of adulthood, respectively) and imaged both groups at A7. The A2 group recapitulated the result from the full turnover experiment, and the A4 group showed a similar amount of turnover as the A2 group, consistent with Model 1 (Figure 4D). We also assessed the presynaptic GFP/RFP ratio in animals that were pulsed at A2, then cultivated 25 °C instead of the standard 20 °C until imaging at A7. Animals age faster, have a shorter lifespan, and exhibit compromised proteostasis at 25 °C; still, the presynaptic SNG-1::ARGO GFP/RFP was about the same (Figure 4D); therefore, these results also align better with Model 1 than Model 2.

### Evidence for endocytic sorting of SNG-1 in presynapse remodeling

*C. elegans* has a series of GABAergic motor neurons with cell bodies along the ventral nerve cord. A subset of these, called the VD neurons, make presynapses along the ventral nerve cord. In the rest, called the DD neurons, presynapses are located along the dorsal nerve cord for most of the animal’s life. Quantifying SNG-1::ARGP GFP/RFP in the VD and DD neurons at different animal ages, we found a strong age-by-neuron-identity interaction in regulating steady-state presynaptic SNG-1::ARGO GFP/RFP (Figure 5). At the L2 larval stage, the VD neurons’ mean ratio was 2.5, whereas the DD neurons’ mean ratio was less than 1.5 (Figure 5A-B). Presynapse size, based on the SNG-1::ARGO RFP intensity, was similar in L2 larvae between the VD and DD neurons (Figure 5C). The DD neurons’ GFP intensity was low in L2 larvae, though (Figure 5D). We speculate that this difference between VD and DD in L2 larvae is related to the distinct mechanisms of synaptic development between these two otherwise similar classes of neuron. In the L1 larval stage, the DD neurons’ presynapses are located along the ventral nerve cord, and the VD neurons have not yet been generated. At the end of the L1 stage, the DD neurons’ presynapses are relocated to the dorsal nerve cord (White *et al*., 1978; Hallam and Jin, 1998). This remodeling involves moving the pre-existing presynaptic proteins from the ventral to the dorsal side (Park *et al*., 2011). Meanwhile, the VD neurons are born, and they generate their initial presynapses along the ventral nerve cord, where they remain. By imaging the VD/DD neurons in the L2 stage, we visualized the synapses shortly after this DD remodeling and VD synaptogenesis event. To fill in the age gap between L2 and A2, we performed a separate experiment in which we assessed steady-state SNG-1::ARGO in the VD versus DD neurons at L3 and A1 (Figure 5E-G). In both neuron classes, the SNG-1::ARGO GFP/RFP ratio is significantly higher at L3 than at A1 (Figure 5E). Considering these results together with the L2 versus A2 experiment suggests that the especially low SNG-1::ARGO ratio in the DD neurons during or directly after presynapse remodeling is a transient condition, and the ratio increases by the next larval stage.

**Figure 5.**
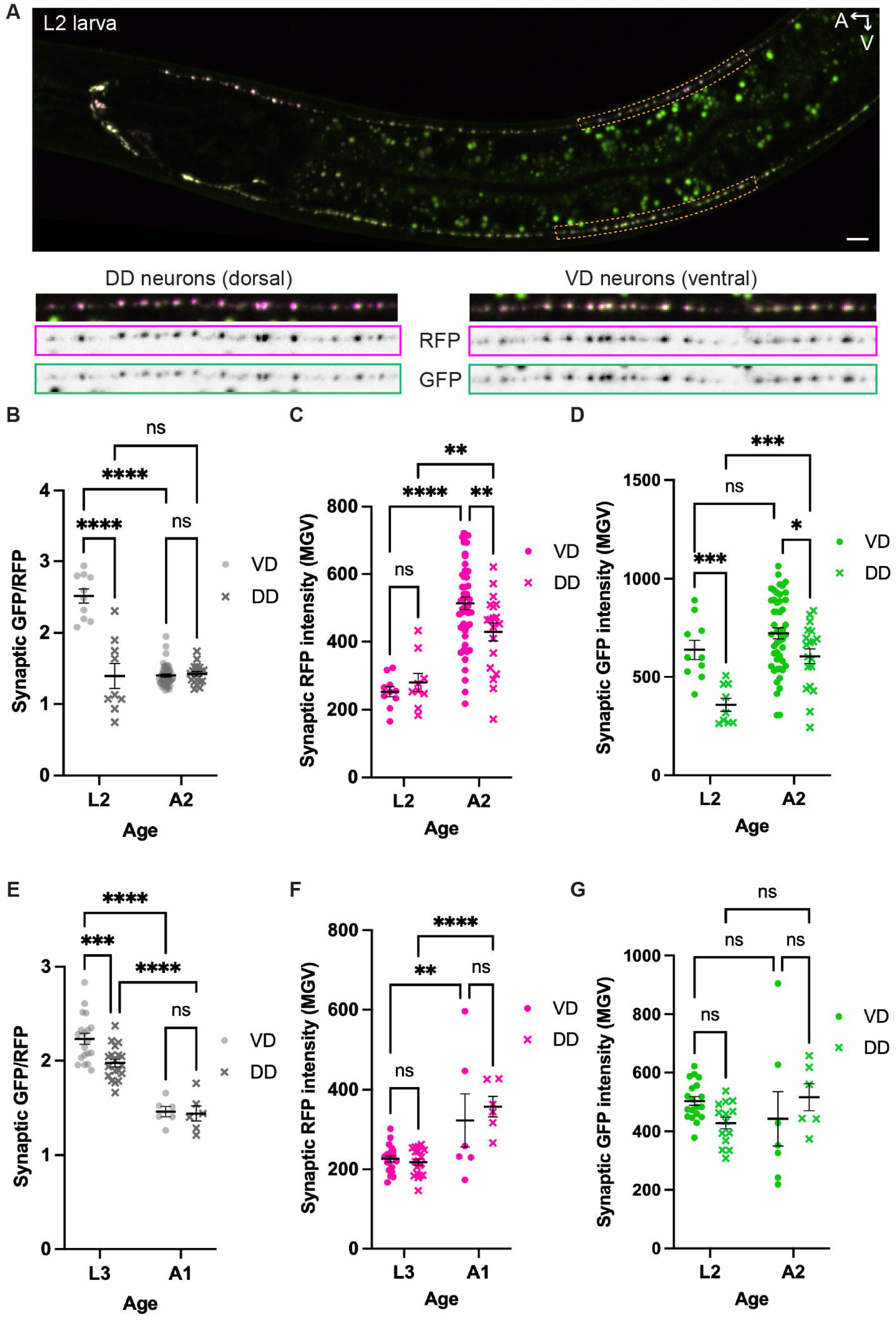
SNG-1::ARGO steady-state GFP/RFP ratio is low in the newly remodeled DD presynapses. (A) Representative image of SNG-1::ARGO in a L2 larva, in which a lower GFP/RFP ratio in the DD neurons compared to the VD neurons is apparent. (B-G) Comparison of presynaptic SNG-1::ARGO GFP/RFP ratio (B,E) and raw fluorescence intensities (C-D,F-G) in the VD versus DD neurons shows a difference at L2 that is abolished by adulthood. Note that the DD ratio at the L2 versus A2 ages is similar (B), whereas the DD ratio at L3 is higher than at A1 (E).Two-way ANOVA with uncorrected Fisher’s LSD, with single pooled variance.

Based on these results, we speculate that when SNG-1::ARGO in the DD neurons is relocated from the ventral to the dorsal nerve cord, it is trafficked within acidified endosomes, wherein the GFP is quenched. Both the SV-transporting kinesin KIF1A/UNC-104 and the more generalist Kinesin-1/UNC-116 are required for DD presynapse remodeling; consistent with our speculation, UNC-116 can transport endosomal and autophagic compartments (Klassen *et al*., 2010; Park *et al*., 2011; Kurup *et al*., 2015; Stavoe *et al*., 2016). In this speculation, the SNG-1::ARGO would be sorted out of the acidified compartments and into SVs in the newly formed DD dorsal presynapses.

## Discussion

In summary, we developed and validated ARGO, a genetically encoded method to easily and inexpensively quantify protein turnover *in vivo* with high spatial and temporal resolution, and we used this method to begin to assess how transmembrane SV protein Synaptogyrin/SNG-1 is turned over from the presynapses in adulthood. These findings provide new insights into SV protein degradation, and they illustrate the utility of the ARGO method.

### The ARGO method

The abundance of each of a cell’s proteins is determined by its rates of production and degradation, plus the rate of cell division (Liu *et al*., 2016). In terminally differentiated cells such as neurons, the protein degradation rate takes a more prominent role in controlling protein abundance (Liu *et al*., 2016). Evidence from proteomic studies indicates that protein half-life can range from minutes to months (and, rarely, even longer), depending on the protein, the cell type, and the cellular context (Toyama *et al*., 2013). An impediment to understanding the regulation of protein degradation is the limits of existing methods for easily assessing protein degradation *in vivo* with high resolution. The most straight-forward existing approach for interrogating a protein-of-interest’s turnover is adding a self-labeling tag to that protein, most commonly SNAP-tag or HaloTag, providing a pulse of fluorescent ligand, then imaging turnover with fluorescence microscopy during the chase (Bojkowska et al. 2011; Bulovaite et al. 2022). ARGO is a conceptually and methodologically similar strategy with several advantages: First, because ARGO is fully genetically encoded, once the strain has been generated, performing experiments is simpler and cheaper. This makes it highly amenable to thorough interrogation of surveillance and degradation mechanisms, for example with forward and reverse genetics. Second, because the imaging is ratiometric, it provides higher resolution of protein half-life, and it is advantageous to use with genetic or circumstantial manipulations that alter the steady-state localization of the protein-of-interest. Third, because the GFP fluorescence is sensitive to acidic environments, analysis of steady-state fluorescence can determine whether and where degradation is mediated via sorting into a lysosomal pathway. The design of ARGO is modular, making it amenable to modification. For instance, though we used heat shock promoter to drive Cre expression here, any temporally inducible transcription system can, in theory, be used to provide the pulse.

### SV protein degradation

Our data indicate that SNG-1 is sorted for degradation at the synapse, then transported to the cell body for degradation in lysosomal compartments. This corroborates the most prevailing, though sometimes disputed, model for the spatial dynamics of SV protein degradation (Jin et al. 2018; Winckler et al. 2018).

We found that the rate of SNG-1 turnover is similar across all the *en passant* presynapses within a neuron regardless of their position relative to the cell body (Figure 3C, S2J). This contrasts with post-synaptic protein PSD95, for which the turnover rate differs between post-synapses within a single neuron based on their spatial positioning in the mouse brain (Bulovaite et al. 2022). Perhaps the continuous trafficking of SVs between synapses generates the conditions in which SNG-1 functions as a single super-pool in regard to turnover (Staras et al. 2010; Herzog et al. 2011; Wu et al. 2013). Alternatively, the molecules and mechanisms that regulate the rate of SNG-1 degradation could be effectively uniform across presynapses.

Our results indicate that reduction-of-function of the sole *C. elegans* E1 ubiquitin ligase, *uba-1*, leads to reduced steady-state presynaptic SNG-1 abundance as well as dramatically slowed SNG-1 turnover from the presynapse. We have recently shown that loss of Transcription Factor EB/HLH-30 likewise causes reduced steady-state presynaptic SNG-1 abundance along with slowed presynaptic SNG-1 turnover . These two phenotypes together suggest that there may be homeostatic feedback that adjusts the steady-state abundance based on the turnover rate or vice versa. On the other hand, these genetic manipulations could cause reduced presynaptic abundance of SNG-1 indirectly through regulation of other proteins within the neuron and/or in other cell types within the animal.

The steady-state presynaptic SNG-1::ARGO GFP/RFP ratio varies with both neuron identity and age. We interpret this variation primarily as reflecting differences in the proportion of SNG-1::ARGO residing in acidic compartments because (A) sorting into acidic compartments is the canonical route for transmembrane protein degradation, (B) the ratio increases upon injection with Bafilomycin A (Figure 2E) and concanamycin A (Zhong and Richardson, 2025), and (C) the marked difference in presynaptic ratio between VD and DD neurons in L2 larvae is not readily explained with alternate, technical interpretations (Figure 5). Still, we cannot exclude that some factor might contribute to changes in SNG-1::ARGO ratio, such as misfolding of GFP on a subset of newly synthesized SNG-1::ARGO molecules. Importantly, the quantification of SNG-1::ARGO half-life normalizes against the GFP/RFP ratio in age-matched control animals and is therefore unaffected by the variation in steady-state ratio.

Positing that the SNG-1::ARGO GFP/RFP ratio mainly reflects sorting into acidic compartments, our data indicate that a large proportion of presynaptic SNG-1 resides in acidic compartments in adult neurons and in newly remodeled DD synapses. These acidic compartments are likely late endosomes and/or autophagosomes. Have these SNG-1 molecules been decommissioned and committed to degradation? This notion seems surprising, yet it aligns with findings from cultured hippocampal neurons showing that SV proteins participate in SV cycling for less than half of their lifetime and, once targeted for degradation, often continue to reside at the presynapse (Truckenbrodt *et al*., 2018). Moreover, sorting into acidic endosomal compartments need not entail an irreversible exit from the SV cycle: intraluminal vesicles within late endosomes can undergo back-fusion, allowing resident proteins to rejoin the functional pool (Eden and Futter, 2021; Perrin *et al*., 2021). It is also possible that SNG-1 has an unappreciated function within endosomal compartments. For the DD synapses in L2 larvae in particular, it would be puzzling if SNG-1 were relocated from the original ventral presynapses to the newly generated dorsal presynapses only to be rapidly eliminated.

Finally, our data suggest that in at least one *C. elegans* neuron type – the GABAergic motor neuron VD2 – SNG-1 exists in two distinct pools within each presynapse that do not intermix over at least several days. This is surprising because SV cycling is generally thought to intermix vesicle proteins in an endosomal sorting compartment, effectively generating a single pool of SV protein over long timescales. We consider several ways two SNG-1 half-lives could arise. One possibility is that SNG-1 resides on two distinct SV pools in the VD2 neuron. To remain chronically distinct, one pool might never participate in neurotransmitter release, the pools might operate in temporally or spatially distinct SV cycling, or one or both pools might exclusively use kiss-and-run fusion, bypassing endosomal sorting.

Hints from other experimental animals suggest these scenarios might be plausible. For instance, a synapse can contain a “resting” SV pool that is resistant to recruitment upon strong experimental stimulation, might not cycle under some physiological contexts, and can comprise over half of the SVs in a presynapse (Chanaday *et al*., 2019). SVs that do release can be further separated into the evoked release versus the spontaneously released pools, and in some contexts, these pools remain segregated for tens of minutes (Sara *et al*., 2005; Mathew *et al*., 2008; Andreae *et al*., 2012; Kavalali, 2014; Egashira *et al*., 2022). Nonetheless, we are not aware of examples where SV pools retain their distinct identities for days. Another possible explanation is that SNG-1 could itself exist in two distinct forms, such as differential post-translational modification(s) during SNG-1 synthesis or a developmental time-window, with one form resistant to degradation. Various mammalian synaptogyrin isoforms can carry post-translational modifications, though none have been identified for *C. elegans* SNG-1 (iPTMnet)(Huang *et al*., 2018). Further research on this two-phase SNG-1 turnover will illuminate how SV proteins can achieve selective turnover, which could have broad implications for how neurotransmission is tuned.

## Materials and Methods

### *C. elegans* culture and maintenance

Nematodes were grown at room temperature on nematode growth media (NGM) plates seeded with *Escherichia coli* OP50. All experiments used hermaphrodite worms and the stages used for each experiment are outlined in each figure legend. Several strains were provided by the Caenorhabditis Genetics Center (CGC), which is funded by NIH Office of Research Infrastructure Program (P40 OD010440). The *carIs1(Pglr-4>Flippase)* allele, which flips-on *sng-1::argo* in the DA9 neuron, was generated by integrating *shyEx246* from the Shaul Yogev lab. To induce expression of *heSi160[Pheat-shock>Cre]* for the ARGO pulse, worm plates were shifted to a 34 °C incubator for 1 hour. For experiments with *uba-1(it129ts)*, animals were grown at 20 °C, then shifted to 25 °C, the non-permissive temperature, at the L2 larval stage. Consistent with reported phenotypes, these animals at 25 °C were fully sterile. The canonical permissive temperature for *uba-1(it129ts)* is 15 °C, where the allele behaves like a weaker hypomorph; in our hands, the strain is viable and appears grossly wild-type at 20 °C.

<u>Strains used in this study:</u>

PBT68 *bqSi506(Prgef-1>Flp, unc-119(+)) IV; sng-1(syb3140car2) heSi160 X*

PBT67 *carIs1(Pglr-4>Flp, Podr-1>rfp) II; sng-1(syb3140car2) heSi160 X*

PBT188 *bqSi488(Ptph-1>Flp, unc-119(+) IV; sng-1(syb3140car2) heSi160 X*

PBT231 *bqSi542 IV; sng-1(syb3140car2) heSi160 X*

PBT84 *sng-1(syb3140car2)*

RB503 *sng-1(ok234) X*

PBT72 *carIs1 II; sng-1(syb3140car2) X*

PBT177 *carIs1 II; uba-1(it129) IV; sng-1(syb3140car2) heSi160 X*

PBT58 *bqSi542(Punc-47>Flp, unc-119(+) IV; sng-1(syb3140) heSi160 X*

### Genome editing using CRISPR/Cas9

The initial ARGO allele, *sng-1(syb3140)*, consisting of *FRT-stop+3’utr-FRT-RFP-loxP-GFP-loxP,* was generated by SunyBiotech. We determined that the resulting SNG-1::RFP::GFP was destabilized and had accelerated degradation. Removal of GFP upon ARGO activation led to increased RFP fluorescence intensity up to the expected intensity based on existing strains (Schwartz and Jorgensen, 2016), indicating that the amino acid linker between the RFP and GFP was causing the destabilization. We therefore used CRISPR/Cas9 to modify the amino acid linker between the RFP and GFP in the *sng-1(syb3140)* allele, generating the *sng-1(syb3140car2)* allele, which fixed the destabilization. The *syb3140car2* allele was used in all experiments in this study with two exceptions: Fig S1M, which quantifies the rate of Cre-mediated recombination, and Table 1, which quantifies Cre/loxP specificity and efficacy. Both of these parameters are expected to be identical between *syb3140* and *syb3140car2*. The *car2* allele was generated following the protocol in (Paix *et al*., 2015). A ssDNA with 36 and 38 bp homology on either side was used for the repair template (Ultramer, Integrated DNA Technologies (IDT)). Two cRNAs (IDT) were used, one on either end of the edit. The genomic sequence of the *argo* cassette and the cDNA sequences of *sng-1::argo* before and after the removal of *GFP* are provided Figures S4-6.

### Aldicarb assay

Aldicarb assays were performed as described (Mahoney et al., 2006). Briefly, 30 Day 1 adults of each strain were placed on a plate containing 0.5 mM aldicarb, and they were periodically assessed for paralysis visually after touch with an eyelash. Plates were scored blind to genotype, and the experiment was executed in three biological replicates, each on a separate day.

### qPCR of genomic DNA

Day 0/L4 animals of the genotype *bqSi542; sng-1(syb3140) heSi160* were shifted to 34 °C for 1 hour at Time = 0. Directly before the shift, and at the indicated times afterward, approximately 50 animals were picked and lysed with the standard worm lysis protocol. Relative abundance of the Cre-recombined locus was assess using primers that aligned to the 3’ end of *RFP* before the first *loxP* and the 3’ end of *sng-1* after the second *loxP.* Relative transcript abundance was calculated using the ΔΔCt method, with primers against *gpi-1* used as the internal control. All reactions were performed in triplicate.

### RNA extraction and qPCR

Total RNA was extracted from *Caenorhabditis elegans* samples using the TRI reagent (Sigma Aldrich), ethanol precipitation protocol. The LunaScript Kit (NEB) was used for complementary DNA (cDNA) synthesis according to the manufacturer’s protocol. Quantitative RT-PCR was performed using the Roche Lightcycler 480. Primers for the target genes (*sng-1, hsp-16*) and the reference/normalization gene (*pmp-42*) were validated empirically for specificity and efficiency. Melting curve analysis was performed at the end of each run to confirm amplification specificity. Gene expression levels were calculated using the ΔΔCt method, with the *pmp-42* gene serving as the internal control. All reactions were performed in triplicate.

### Bafilomycin A treatment

Bafilomycin A was injected into the body cavity of *C. elegans* similarly to previous work (Wilkinson et al., 2015). Briefly, 25 μM of BafA in 5% DMSO or 0.2% DMSO (“mock”) was co-injected with Orange G for color into the pseudocoelom, and animals were allowed to recover for three hours before they were imaged with fluorescence confocal microscopy as described above. BafA injected animals were still moving when they were put on slides, indicating that the BafA had not completely blocked V-ATPase function.

### Assessing specificity and efficacy of the pulse

To assess the efficacy and specificity of Heat-shock>Cre-mediated recombination of *GFP* from the ARGO cassette (Table 1), fluorescence was scored based on visualizing SNG-1::ARGO GFP and RFP fluorescence at least 3 days after activation. Scoring was performed using a combination of two approaches: 1) qualitative assessment by eye on a compound fluorescent microscope, wherein the experimenter performed an initial visual assessment of a subset of animals and then binned all of the animals in the cohort as either “bright” (no recombination) or “dim” (recombination); 2) semi-quantitative assessment on the spinning disk confocal microscope, wherein the animals were imaged with standardized microscope settings, raw fluorescence intensity and GFP/RFP values were assessed for a subset of animals, and then all animals in the cohort were binned as “ratio high” (no recombination) or “ratio low”(recombination). The distribution of GFP/RFP ratios in all instances was bimodal; if an animal appeared to have an intermediate value, which occurred in about 1-5% of individuals, that animal was not scored.

### Microscopy and imaging

Worms were mounted on 3% freshly made agarose pads into drops of 20 mM levamisole. An imaging system comprised of a Nikon Eclipse Ti2 microscope, Yokogawa CSU-W1 SoRa spinning-disk unit and a Hamamatsu ORCA-fusionBT digital camera C15440 with Plan Apo x60 A1/2 Nikon objective with water immersion was used to acquire all images. Image settings (60% of laser power and 200 ms exposure time) were identical for all groups and treatments across experiments. Z-stacks were taken with 0.4 μm step in the 20-30 μm range, depending on type of neuron. For super resolution by optical reassignment (Figure 1I), the 2.8x SoRa magnifier was used, z-stacks were acquired with 0.3 μm steps, and images were deconvoluted in post-processing.

### Microscopy data analysis

Nd2 files were screened with NIS Elements Software for unusually low signal, severe movement or drifting and discarded, the rest images were processed with Fiji (ImageJ) software. Image stacks were transformed using z-projection function with max intensity option, areas of interest were selected either with Rectangle tool (for cell bodies) or Segmented line and Straighten function (for synaptic areas). For each image, a Threshold was applied with default settings to the RFP channel, and the Analyze Particles function was used to identify ROIs in that thresholded image. These ROIs were then used to quantify fluorescence from both the RFP and GFP channels. The mean intensity of each channel was used for all calculations. Each obtained array of values in csv format was united in a single data frame, and synaptic puncta less than 0.1 µm² (for the wild-type DA9 and NSM datasets) or 0.2 µm² (for the VD, DD, and *uba-1* mutant datasets) were filtered out due to high variability in relative RFP:GFP fluorescence intensities. This high variability was likely due to low signal-to-noise; a major contributing factor to the noise was autofluorescence. The GFP/RFP ratio calculation was done for each presynapse with Jupyter Notebook and Python scripts.

To find the average presynaptic GFP/RFP ratio per neuron, all the individual GFP/RFP ratios within the presynaptic region of a neuron were averaged.

To find the average presynaptic GFP/RFP ratio by position along the proximal-distal axis, the length of each neuron’s presynaptic region was divided into the most proximal 30%, the most distal 30% and the middle 40%, and all the presynaptic puncta within each domain was averaged.

### Statistical Analyses

Prism GraphPad software was used to generate all plots and perform most statistical analyses.

The rest of the statistical analyses were performed in Rstudio as noted below.

### Assessing the effective timing of the pulse from microscopy data

To test whether the SNG-1::ARGO GFP/RFP decreased significantly by 4 hours after the heat-shock (Figure S1F-L), we used a linear mixed-effects model with treatment (no heat-shock vs 4 hours post heat-shock) as a fixed effect and worm as a random effect to include the GFP/RFP ratio measurements for each presynapse and account for multiple measurements per animal (lmer function, lme4 package). Residual diagnostics were assessed using the DHARMa package. For DA9 D2 and D6 (Figure S1H, J), data were log transformed to meet model assumptions, and the back-transformed mean GFP/RFP per animal are plotted. For DA9 D2 with *heSi160* (Figure S1H), residual plots with the log transform still showed a deviation from normality (KS test: ns, Dispersion test: ns, outlier test, p = 0.012), but model assumptions were considered adequately met given the large sample size and consistency of results across multiple statistical approaches, including nonparametric tests. For NSM D2 (Figure S1K), data were square-root transformed to meet model assumptions, and the back-transformed mean GFP/RFP per animal are plotted. Empirically, this transformation was the closest to fitting model assumptions, though residual plots still showed some deviation from normality (KS test: ns, Dispersion test: ns, outlier test, p = 0.016). For VD2 D2 (Figure S1L), data did not fit the assumptions of the linear mixed model and an adequate transformation could not be found, so the mean SNG-1 ARGO GFP/RFP ratio per animal was analyzed using a two-sided t test (the data distribution was not significantly different from a normal distribution).

As another approach to assess whether and to what extent a cell sometimes ectopically recombines one of the two alleles of *sng-1::argo*, we compared the average presynaptic SNG-1::ARGO GFP/RFP mean and standard deviation in the DA9 neuron between a strain with vs without *heSi160[Pheat-shock>Cre]* and saw no significant difference (Figure S1A).

As a third approach to assess specificity and efficacy of recombination, we compared the interneuronal coefficient of variation (CV) in the presynaptic SNG-1::ARGO GFP/RFP between control versus pulsed groups. We calculated the CV for control versus heat-shocked groups at each timepoint for each neuron class. As there is one CV value per neuron class, treatment, and timepoint, these comparisons are qualitative. If the CV is roughly constant in the non-pulsed group over time, it would suggest that there is not much ectopic recombination without the pulse treatment. Similarly, if the CV is roughly constant in the pulsed group over time, it would suggest that recombination is uniformly completed in both alleles of each animal. *I haven’t added this yet. It would be most useful to include the version with all data vs after outliers are removed. Is this necessary?*

### Assessing Variance

To assess variability in SNG-1::ARGO GFP/RFP ratios across synapses within individual neurons (intraneuronal variance), because the data roughly followed a lognormal distribution, we used Rstudio to calculate the CV as:

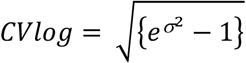

Then, average of all CVs per worm was compared between days using Kruskal-Wallis test, because the data were nonparametric, with Dunn’s multiple comparisons test.

### Fitting decay curves to turnover data

Each SNG-1::ARGO GFP/RFP datapoint from the turnover experiments is the mean of all the GFP/RFP ratios from each presynaptic puncta detected in a single animal. Those datapoints were normalized by the mean presynaptic GFP/RFP ratio from an age-matched control (non-pulsed) cohort of animals that was imaged in parallel. In this way, the steady-state changes in presynaptic SNG-1::ARGO GFP/RFP ratio by animal age do not impact the turnover calculations. SNG-1::ARGO half-life was calculated using Prism to fit a one-phase exponential decay curve to the data. The curves were fitted using least squares regression with no weighting, considering each replicate Y value as an individual point. Outliers were detected and eliminated with Q = 5% (DA9), 1% (NSM), 0r 3% (VD2). These values were selected based on the data in Table 1, which assessed the frequency of ectopic *gfp* excision in the absence of the heat-shock pulse, and the frequency of failed *GFP* excision following the heat-shock pulse, for each neuron identity. Unless otherwise noted, we constrained the plateau to the experimentally determined value based on background GFP fluorescence, which comes from autofluorescence. This was measured from the adult Day 7 images for each of the three neuron identities by hand-selecting the synaptic region in the RFP channel, copying it to the GFP channel, shifting the selection to a position adjacent to the synapses, then finding the GFP fluorescence intensities inside the ROIs. Goodness of fit was quantified with R squared.

### Assessing VD2 turnover

We noticed that for the VD2 neuron, the presynaptic SNG-1::ARGO GFP/RFP ratio does not appear to approach the background in the turnover experiment. Indeed, a two-phase exponential decay function improves the curve fit, from an R squared of 0.41 to 0.48, and comparing the one-phase versus two-phase functions using extra sum-of-squares F test showed that the two-phase curve was significantly favored (Null hypothesis: one phase decay, alternative hypothesis: two phase decay, P = 0.0175). A two-phase exponential decay function will often produce a better fit than a one-phase function due to its additional parameters. Therefore, we took a second approach to assess whether the one-phase decay function could be ruled out using the same dataset: we compared the predicted plateau value from the one-phase function versus the measured background. Specifically, we fit a one-phase exponential decay function to the data without constraining the plateau. We compared the best-fit plateau and 95% C.I. from that function to the mean background and 95% C.I. calculated from the microscopy images (as described above)(Figure 4B). Indeed, the one-phase curve plateaus above the background, and the 95% C.I.s do not overlap (Figure 4B). Furthermore, in Figure 4D, we use the best-fit parameters from the two-phase exponential decay function from the VD2 turnover dataset in Figure 4A to predict the presynaptic GFP/RFP ratio 5 days after the pulse (the Model 1, D2 bar). That predicted value matched the real value ± SEM from a separate experiment, showing that this result is robust (the Results A2 bar).

Given that we could not find evidence for different turnover rates between individual presynaptic puncta within the VD2 neuron (Figure 4C), we considered two explanations for the two-phase turnover kinetics of SNG-1::ARGO within individual VD2 presynapses:

Model 1: there are two chronically distinct pools of SNG-1 protein

Model 2: there is a time-dependent suppression of turnover

We generated predictions from the two proposed models as described below. Then, we compared those predictions qualitatively to an experiment in which we pulsed the VD2 SNG-1::ARGO at adult D2 versus D4 of adulthood and imaged at D7 (Figure 4D). In order to prevent datapoints from inefficient or spontaneous Cre-mediated recombination from skewing the results, we censored aberrant data points as follows: the median presynaptic GFP/RFP ratio was calculated for each set of A7 animals imaged (one data point per animal, which was the mean GFP/RFP ratio across all VD2 presynaptic puncta for that animal): control, pulse at A2, pulse at A4 (all incubated at 20 °C), and control versus pulse at A2 and incubated at 25 °C. For the A7 controls, datapoints lower than the median ratio for the pulse at A4 group were censored (3 of 51 for the 20 °C controls, 0 of 20 for the 25 °C controls). For all three pulsed groups, datapoints higher than the median ratio for the control A7 group at the corresponding temperature were censored (6 of 44, 13.6% for A2 pulse; 6 of 19 for the A4 pulse; 3 of 29 for the A2 pulse and 25 °C incubation).

#### I. Assessing Model I: that there are two distinct populations of SNG-1 with different turnover kinetics at each synapse

Using the best-fit values from the two-phase exponential decay function from the data in Figure 4D, we predicted the presynaptic GFP/RFP ratio 3 days and 5 days after the pulse using ChatGTP 5.

#### II. Assessing Model 2: that turnover is suppressed during the course of the turnover experiment

In this model, all presynaptic SNG-1::ARGO is initially subject to fast-phase turnover, but this turnover becomes inhibited or dysfunctional at a specific time in the animal’s life, which occurs during the course of the turnover experiment when the pulse is at A2. In this model, only protein that undergoes fast-phase turnover prior to the onset of inhibition is degraded; all remaining protein is retained with unappreciable turnover thereafter. In other words, the final GFP/RFP ratio reflects the amount of SNG-1::ARGO that experienced degradation before the inhibition/dysfunction event. For the fast-phase, k = 0.7535 day^-1^. Solving for the time at which 60% of the protein remains gives 0.66 d after the pulse, which would equate to A2.66.

## Data availability

Strains and plasmids are available upon request. Microscopy images will be deposited in BioImage Archive.

## Acknowledgments

Some strains were provided by the *Caenorhabditis* elegans Genetics Center (CGC), which is funded by the National Institutes of Health (NIH) Office of Research Infrastructure Programs (P40 OD010440). The Shaul Yogev lab provided *shyEx246*, which was integrated to generate *carIs1*. We thank Michael O. Harding from the Statistical Consulting group at the University of Wisconsin – Madison for assistance in data analysis. We thank Julianna Miller, Emma Keep, Adam Darlington, Margaret Brener, and Sophia Whitley for assistance in the lab. We thank Ruben H. Land and David M. Lipton for feedback on the manuscript. This work was supported by the National Institutes of Health (NIH) grant R35 GM154869 to C.E.R.

