## Supplemental Figures 1-6 for "Analysis *in vivo* using a new method, ARGO (Analysis of Red Green Offset), reveals complexity and cell-type specificity in presynaptic turnover of synaptic vesicle protein Synaptogyrin/SNG-1"

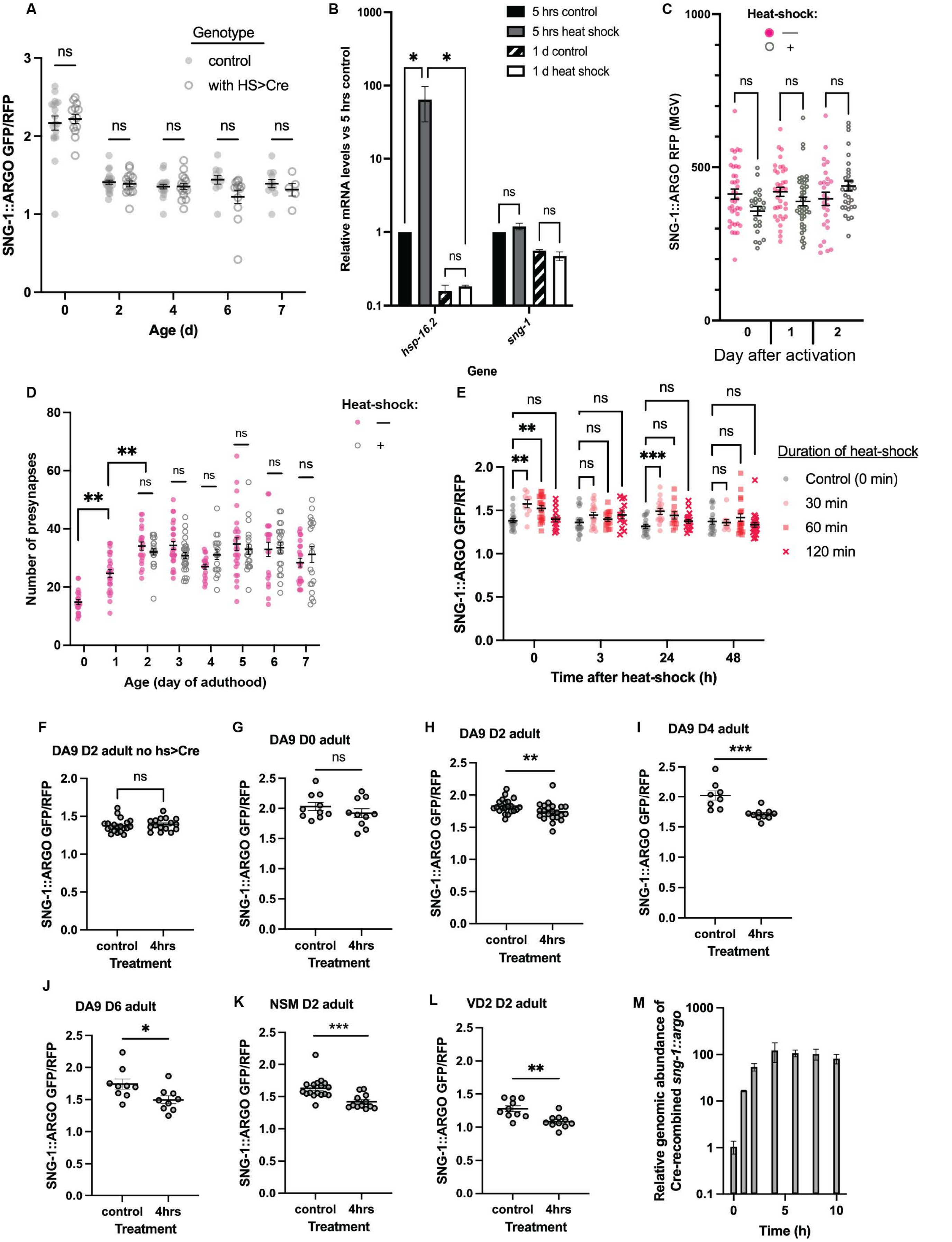


Figure S1. Validation of the heat-shock-generated pulse to express Cre recombinase. (A) Steady-state average presynaptic SNG-1::ARGO GFP/RFP in a strain without versus with the Pheat-shock>Cre transgene heSi160 indicates that there is little ectopic recombination in the absence of heat-shock. Each data point is the average of all individual presynaptic ratios from one DA9 neuron. (B) The heat-shock pulse does not discernably impact sng-1 transcript levels. qRT-PCR analysis of sng-1 mRNA levels with versus without the heat-shock conditions used as the pulse to activate SNG-1::ARGO. Values represent fold change relative to control (no heat-shock) ± SEM (n = 2 biological replicates, *P<0.05, two-way ANOVA with Tukey post-test). (C-D) The heat-shock pulse shows no significant effect on SNG-1 ARGO steady-state abundance. (C) Mean presynaptic SNG-1::ARGO RFP fluorescence intensity with versus without the heat-shock pulse. The experiment was started on A2. Each data point shows the average mean RFP intensity for all synapses of the DA9 neuron for one neuron. (D) Number of DA9 presynapses per neuron with versus without the heat shock pulse at A2. For (C-D), The heat-shocked animals at time = 0 were imaged 4 hours post heat-shock, and bars show mean ± SEM (ns (not significant): P>0.05, *P<0.5, ***P<0.001, two-way ANOVA with Tukey post-test). (E) Quantification of presynaptic SNG-1::ARGO GFP/RFP in the DA9 neuron, in a strain lacking heSi160[Pheat-shock>Cre]. Animals were heat-shocked at 34 ⍰C for the indicated time, or not at all, at adult Day 2, then periodically imaged. Each data point is average per worm; bars show mean ± SEM (ns (ns: P>0.5, **P<0.01, ***P<0.001, two-way ANOVA with Tukey post-test. The heat-shock used for experiments throughout the manuscript was 34 ⍰C for one hour. (F-L) A drop in the presynaptic SNG-1::ARGO GFP/RFP can typically be detected by 4 hours post heat-shock pulse in the presence of the Pheat-shock>Cre transgene (G-L), but not in its absence (F). Analyses are described in Materials and Methods. The data used in F are a subset of the data from E. (M) Assessing the rate of genomic recombination to remove the gfp after the pulse, applied to Day 0 adults via qPCR. Time = 0 was immediately after the 1-hour heat shock (of note, some recombination is expected to have occurred already due to the quick activation of HSF-1 to drive Phest-shock>Cre expression upon the initiation of heat-shock).

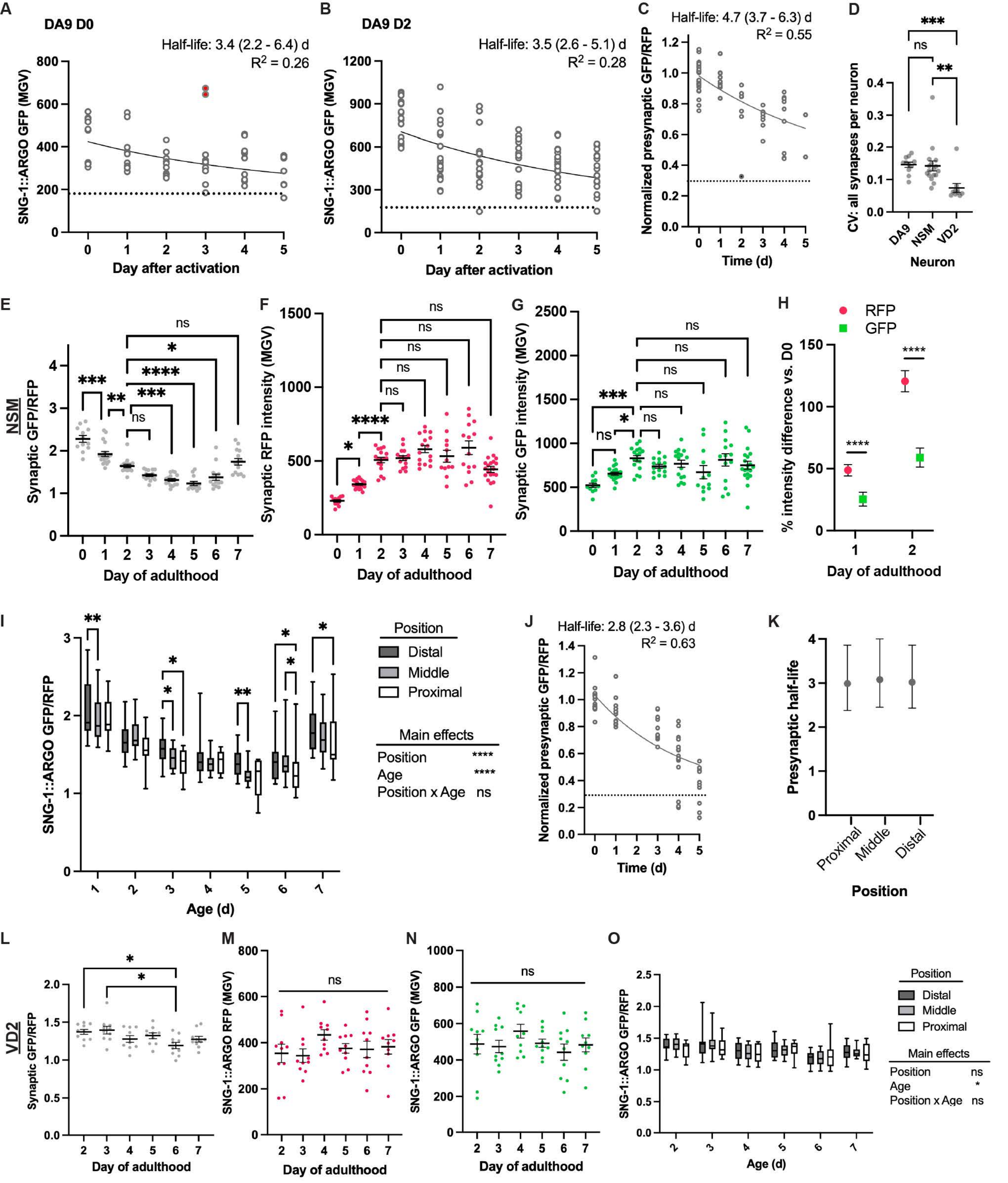


Figure S2. Additional analyses of SNG-1::ARGO steady-state fluorescence and turnover. (A-B) Quantification of SNG-1::ARGO degradation in the DA9 neuron using the mean GFP intensity at each presynapse. Each data point is the average of all the mean GFP values for one animal. Data were fit to one-phase exponential decay curves with the plateau set to the experimentally calculated plateau, which arose from autofluorescence in the GFP channel. Filled circles indicate outlier data points that were excluded from the analysis. The data for these graphs came from the same experiments as those used to generate Figures 3B and 3E. Note that for the pulse at A0, the calculated SNG-1 half-life is longer using mean GFP, which only quantifies degradation, compared to using the GFP/RFP ratio (Figure 3E), which quantifies degradation plus synapse growth. By contrast, with the pulse at A2, the half-life calculated using mean GFP is indistinguishable from that calculated using the GFP/RFP ratio, though the 95% C.I. is substantially larger. (C) A turnover experiment in the DA9 neuron with the pulse at A2, wherein the animals were maintained at 25 C after the pulse. These results are included for comparison to the uba-1(it129ts) turnover experiment; however, note that this is not the ideal comparison because the uba-1(it129ts) mutants were maintained at 25 C from the L2 larval stage. (D) Comparison of intraneuronal variance between GFP/RFP ratio across individual presynapses. Each data point is the CV for all the presynaptic puncta in one neuron. (E-G) Average presynaptic RFP/GFP ratio (E), and the underlying intensity data for RFP (F) and GFP (G) from SNG-1::ARGO in the NSM neuron. Data points are each the average of all presynaptic puncta from a single neuron; bar shows mean ± SEM. (H) Percent change in average presynaptic RFP and GFP intensity from A0 through A2. Plotted is mean ± SEM calculated from the data in F-G. (I) Steady-state SNG-1::ARGO presynaptic GFP/RFP by presynapse position along the axon in the NSM neuron. (J) One-phase exponential decay function fit to the data from a D2 SNG-1::ARGO turnover experiment in the NSM neuron; each data point shows the average presynaptic SNG-1::ARGO GFP/RFP ratio from all the presynpases in a single neuron. Dashed black line shows the experimentally calculated background, which comes from autofluorescence. Filled circles show outlier data points that were excluded from the curve calculation. (K) SNG-1::ARGO presynaptic half-life by proximal-distal position relative to the NSM neuron cell body (mean ± 95% C.I.). (L-N) Average presynaptic RFP/GFP ratio (L), and the underlying intensity data for RFP (M) and GFP (N) from SNG-1::ARGO in the VD2 neuron. Data points are each the average of all presynaptic puncta from a single neuron; bar shows mean ± SEM. (O) Steady-state SNG-1::ARGO presynaptic GFP/RFP by presynapse position along the axon in the VD2 neuron. ns: not significant, *P<0.05, **P<0.01, ***P<0.001, ****P<0.0001, one-way ANOVA or two-way ANOVA with Tukey post-test. For H and N, comparisons that were not significant are not shown.

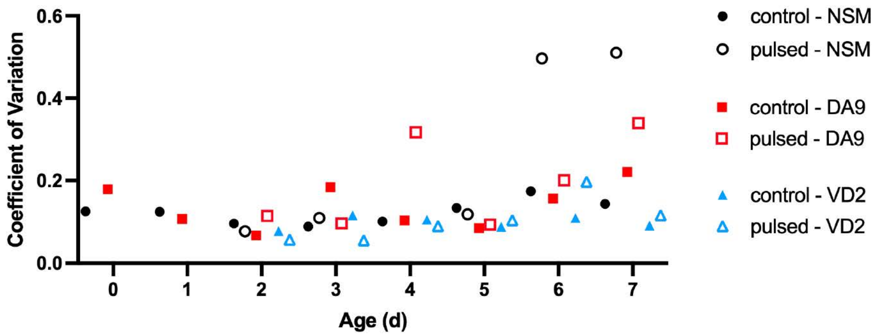


Figure S3. Interneuronal Coefficient of Variation (CV) for each neuron identity during steady-state imaging and the turnover experiment. The analyses were performed on the same datasets used throughout the manuscript for steady-state imaging and turnover with the pulse at A2. As there is one data set per neuron identity, there is one CV per neuron/age/treatment. This is therefore a qualitative assessment of SNG-1::ARGO specificity and efficacy by neuron. It complements the results in Table 1 in that it is better-designed to detect instances wherein a single allele of gfp is removed. Note that the VD2 neuron does not show higher CVs at any age compared to the DA9 and NSM neurons, so the two-phase turnover in the VD2 neuron is not due to unselective or ineffective gfp removal.

(from 1-415 bp)

### cDNA sng-1 syb3140car2 (1619 bp)

```
CCATGCAACAACCACCATCAAACCCATAtACTCAaTCgGAAGGATATGGTTATGGAAGTTCCTATTCTCTAGAAAGTATAGGA
GGTACGTTGTTGGTGGTAGTTTGGGTATaTGAGTtAGcCTTCCTATACCAATACCTTCAAGGATAAGAGATCTTTCATATCCT
     2    4    6    8    10   12   14   16   18   20   22   24   26
 M  Q  Q  P  P  S  N  P  Y  T  Q  S  E  G  Y  G  Y  G  S  S  Y  S  L  E  S  I  G
sng-1:Argo
sng-1 C-term                                   FRT
10   20   30   40   50   60   70   80
```

```
ACTTCAGTCTCCAAGGGAGAGGAGCTCATCAAGGAGAACATGCACATGAAGCTCTACATGGAGGGAACCGTCAACAACCACCA
TGAAGTCAGAGGTTCCCTCTCCTCGAGTAGTTCCTCTTGTACGTGTACTTCGAGATGTACCTCCCTTGGCAGTTGTTGGTGGT
28   30   32   34   36   38   40   42   44   46   48   50   52   54
 T  S  V  S  K  G  E  E  L  I  K  E  N  M  H  M  K  L  Y  M  E  G  T  V  N  N  H  H
sng-1:Argo
tagRFP
90   100   110   120   130   140   150   160
```

```
CTTCAAGTGCACCTCCGAGGGAGAGGGAAAGCCATACGAGGGAACCCAAACCATGCGTATCAAGTCGTCGAGGGAGGACCACT
GAAGTTCACGTGGAGGCTCCCTCTCCCTTTCGGTATGCTCCCTTGGGTTTGGTACGCATAGTTCAGCAGCTCCCTCCTGGTGA
56   58   60   62   64   66   68   70   72   74   76   78   80   82
 F  K  C  T  S  E  G  E  G  K  P  Y  E  G  T  Q  T  M  R  I  K  S  S  R  E  D  H
sng-1:Argo
tagRFP
170   180   190   200   210   220   230   240
```

```
CCCATTCGCCTTCGACATCCTCGCCACCTCCTTCATGTACGGATCCCGTACCTTCATCAACCACACCCAAGGAATCCCAGACT
GGGTAAGCGGAAGCTGTAGGAGCGGTGGAGGAAGTACATGCCTAGGGCATGGAAGTAGTTGGTGTGGGTTCCTTAGGGTCTGA
84   86   88   90   92   94   96   98   100   102   104   106   108   110
 S  H  S  P  S  T  S  S  P  P  P  S  C  T  D  P  V  P  S  S  T  T  P  K  E  S  Q  T
sng-1:Argo
tagRFP
250   260   270   280   290   300   310   320   330
```

```
TCTTCAAGCAATCCTTCCCAGAGGGATTCACCTGGGAGCGTGTCACCACCTACGAGGACGGAGGAGTCCTCACCGCCACCCAA
AGAAGTTCGTTAGGAAGGGTCTCCCTAAGTGGACCCTCGCACAGTGGTGGATGCTCCTGCCTCCTCAGGAGTGGCGGTGGGTT
112   114   116   118   120   122   124   126   128   130   132   134   136   138
 S  S  S  N  P  S  Q  R  D  S  P  G  S  V  S  P  P  T  R  T  E  E  S  S  P  P  P  K
sng-1:Argo
tagRFP
340   350   360   370   380   390   400   410
```

cDNA sng-1 syb3140car2 (1619 bp) (from 416-830 bp)

GACACCTCCCTCCAAGACGGATGCCTCATCTACAACGTCAATCCGTGGAGTCAACTTCCCATCCAACGGACCAGTCATGCAAA
CTGTGGAGGGAGGTTCTGCCTACGGAGTAGATGTTGCAGTTAGGCACCTCAGTTGAAGGGTAGGTTGCCTGGTCAGTACGTTT
140 142 144 146 148 150 152 154 156 158 160 162 164
T P P S K T D A S S T T S I R G V N F P S N G P V M Q
sng-1:Argo
tagRFP
420 430 440 450 460 470 480 490

AGAAGACCCTCGGATGGGAGGCCAACACCGAGATGCTCTACCCAGCCGACGGAGGACTcGAGGGACGTACCGACATGGCCCTC
TCTTCTGGGAGCCTACCCTCCGGTTGTGGCTCTACGAGATGGGTCGGCTGCCTCCTGAgCTCCCTGCATGGCTGTACCGGGAG
166 168 170 172 174 176 178 180 182 184 186 188 190 192
K K T L G W E A N T E M L Y P A D G G L E G R T D M A L
sng-1:Argo
tagRFP
500 510 520 530 540 550 560 570 580

AAGCTCGTCGGAGGAGGACACCTCATCTGCAACTTCAAGACCACCTACCGTTCCAAGCCAGCCAAGAACCTCAAGATGCCAGG
TTCGAGCAGCCTCCTCCTGTGGAGTAGACGTTGAAGTTCTGGTGGATGGCAAGGTTCGGTCGGTTCTTGGAGTTCTACGGTCC
194 196 198 200 202 204 206 208 210 212 214 216 218 220
K L V G G G H L I C N F K T T Y R S K P A K N L K M P G
sng-1:Argo
tagRFP
590 600 610 620 630 640 650 660

AGTCTACTACGTCGACCACCGTCTcGAGCGTATCAAGGAGGCCGACAAGGAGACcTACGTCGAGCAACACGAGGTCGCCGTCG
TCAGATGATGCAGCTGGTGGCAGAgCTCGCATAGTTCCTCCGGCTGTTCCTCTGgATGCAGCTCGTTGTGCTCCAGCGGCAGC
222 224 226 228 230 232 234 236 238 240 242 244 246 248
V Y Y V D H R L E R I K E A D K E T Y V E Q H E V A V
sng-1:Argo
tagRFP
670 680 690 700 710 720 730 740

CCCGTTACTGCGACCTCCCATCCAAGCTCGGACACAAGCTCAACGgTatggaTgaActGtaTaaGGGTTCTGGGAGTGGAAGC
GGGCAATGACGCTGGAGGGTAGGTTCGAGCCTGTGTTCGAGTTGCcAtacctActTgaCatAttCCCAAGACCCTCACCTTCG
250 252 254 256 258 260 262 264 266 268 270 272 274 276
A R Y C D L P S K L G H K L N G M D E L Y K G S G S G S
sng-1:Argo
tagRFP
GSGSGSGS linker
750 760 770 780 790 800 810 820 830

cDNA sng-1 syb3140car2 (1619 bp) (from 831-1245 bp)

```
GGCTCTATGAGTAAAGGAGAAGAACTTTTCACTGGAGTTGTCCCAATTCTTGTTGAATTAGATGGTGATGTTAATGGGCACAA
CCGAGATACTCATTTCCTCTTCTTGAAAAGTGACCTCAACAGGGTTAAGAACAACTTAATCTACCACTACAATTACCCGTGTT
   278   280   282   284   286   288   290   292   294   296   298   300   302   304
G  S  M  S  K  G  E  E  L  F  T  G  V  V  P  I  L  V  E  L  D  G  D  V  N  G  H  K
sng-1:Argo
GFP
      840       850       860       870       880       890       900       910
```

```
ATTTTCTGTCAGTGGAGAGGGTGAAGGTGATGCAACATACGGAAAACTTACCCTTAAATTTATTTGCACTACTGGAAAACTAC
TAAAAGACAGTCACCTCTCCCACTTCCACTACGTTGTATGCCTTTTGAATGGGAATTTAAATAAACGTGATGACCTTTTGATG
    306   308   310   312   314   316   318   320   322   324   326   328   330
 F  S  V  S  G  E  G  E  G  D  A  T  Y  G  K  L  T  L  K  F  I  C  T  T  G  K  L
sng-1:Argo
GFP
   920       930       940       950       960       970       980       990
```

```
CTGTTCCATGGCCAACACTTGTCACTACTTTCTGTTATGGTGTTCAATGCTTCTCGAGATACCCAGATCATATGAAACGGCAT
GACAAGGTACCGGTTGTGAACAGTGATGAAAGACAATACCACAAGTTACGAAGAGCTCTATGGGTCTAGTATACTTTGCCGTA
332   334   336   338   340   342   344   346   348   350   352   354   356   358
 P V  P  W  P  T  L  V  T  T  F  C  Y  G  V  Q  C  F  S  R  Y  P  D  H  M  K  R  H
sng-1:Argo
GFP
  1,000     1,010     1,020     1,030     1,040     1,050     1,060     1,070
```

```
GACTTTTTCAAGAGTGCCATGCCCGAAGGTTATGTACAGGAAAGAACTATATTTTTCAAAGATGACGGGAACTACAAGACACG
CTGAAAAAGTTCTCACGGTACGGGCTTCCAATACATGTCCTTTCTTGATATAAAAAGTTTCTACTGCCCTTGATGTTCTGTGC
360   362   364   366   368   370   372   374   376   378   380   382   384   386
D  F  F  K  S  A  M  P  E  G  Y  V  Q  E  R  T  I  F  F  K  D  D  G  N  Y  K  T  R
sng-1:Argo
GFP
1,080   1,090     1,100     1,110     1,120     1,130     1,140     1,150     1,160
```

```
TGCTGAAGTCAAGTTTGAAGGTGATACCCTTGTTAATAGAATCGAGTTAAAAGGTATTGATTTTAAAGAAGATGGAAACATTC
ACGACTTCAGTTCAAACTTCCACTATGGGAACAATTATCTTAGCTCAATTTTCCATAACTAAAATTTCTTCTACCTTTGTAAG
  388   390   392   394   396   398   400   402   404   406   408   410   412   414
 A  E  V  K  F  E  G  D  T  L  V  N  R  I  E  L  K  G  I  D  F  K  E  D  G  N  I
sng-1:Argo
GFP
     1,170     1,180     1,190     1,200     1,210     1,220     1,230     1,240
```

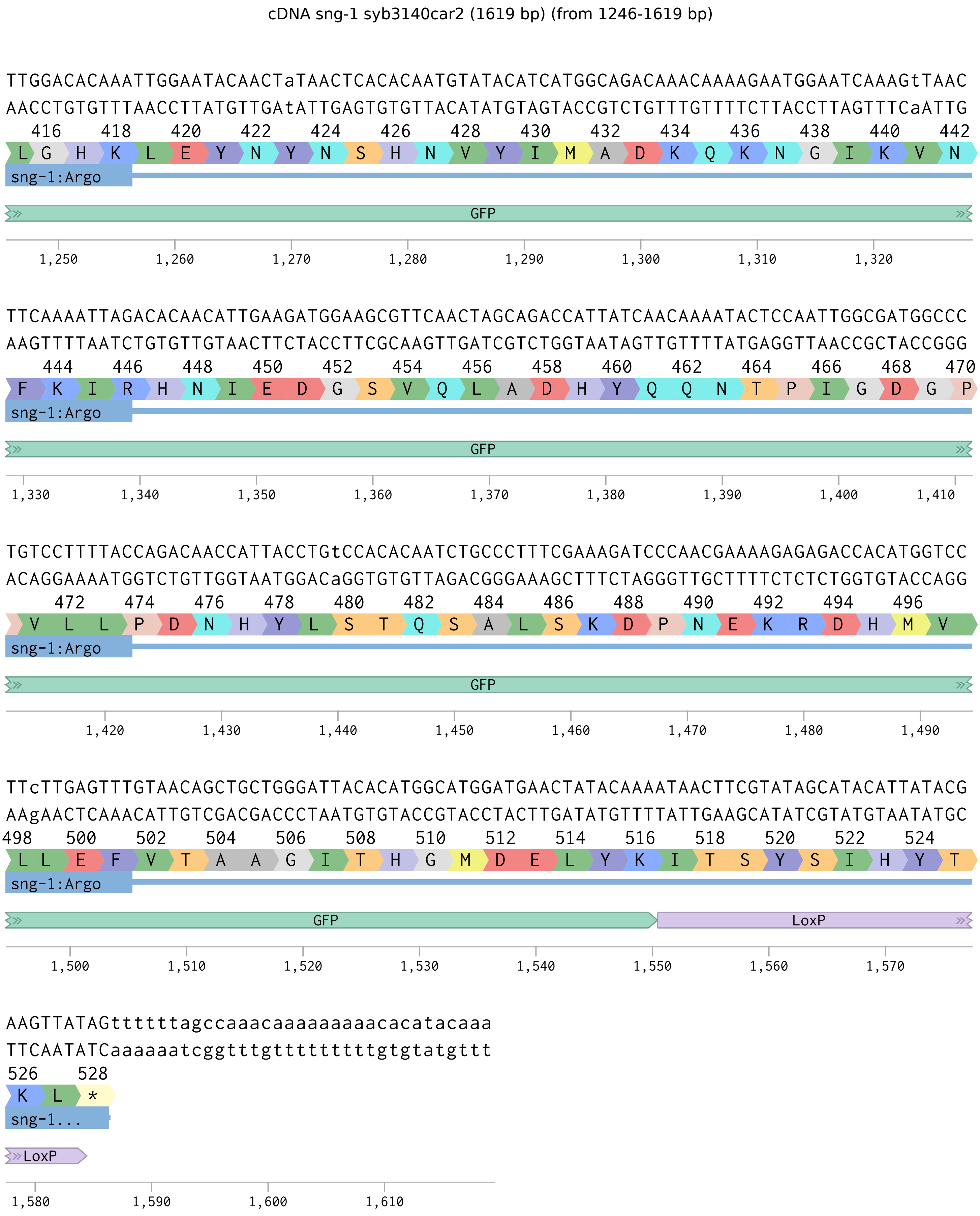


Figure S4. Sequence of sng-1(syb1340car2) cDNA when dually-tagged with RFP and GFP.

(from 1-465 bp)

### syb3140car2 recombined (891 bp)

```
CCATGCAACAACCACCATCAAACCCATAtACTCAaTCgGAAGGATATGGTTATGGAAGTTCCTATTCTCTAGAAAGTATAGGAACTTCAGTCT
GGTACGTTGTTGGTGGTAGTTTGGGTATaTGAGTtAGcCTTCCTATACCAATACCTTCAAGGATAAGAGATCTTTCATATCCTTGAAGTCAGA
  2   4   6   8   10   12   14   16   18   20   22   24   26   28   30
M Q Q P P S N P Y T Q S E G Y G Y G S S Y S L E S I G T S V
Translation
sng-1 C-term    FRT
10 20 30 40 50 60 70 80 90
```

```
CCAAGGGAGAGGAGCTCATCAAGGAGAACATGCACATGAAGCTCTACATGGAGGGAACCGTCAACAACCACCACTTCAAGTGCACCTCCGAGG
GGTTCCCTCTCCTCGAGTAGTTCCTCTTGTACGTGTACTTCGAGATGTACCTCCCTTGGCAGTTGTTGGTGGTGAAGTTCACGTGGAGGCTCC
  32   34   36   38   40   42   44   46   48   50   52   54   56   58   60
S K G E E L I K E N M H M K L Y M E G T V N N H H F K C T S E
Translation
tagRFP
100 110 120 130 140 150 160 170 180
```

```
GAGAGGGAAAGCCATACGAGGGAACCCAAACCATGCGTATCAAGGTCGTCGAGGGAGGACCACTCCCATTCGCCTTCGACATCCTCGCCACCT
CTCTCCCTTTCGGTATGCTCCCTTGGGTTTGGTACGCATAGTTCCAGCAGCTCCCTCCTGGTGAGGGTAAGCGGAAGCTGTAGGAGCGGTGGA
62   64   66   68   70   72   74   76   78   80   82   84   86   88   90   92
G E G K P Y E G T Q T M R I K V V E G G P L P F A F D I L A T
Translation
tagRFP
190 200 210 220 230 240 250 260 270
```

```
CCTTCATGTACGGATCCCGTACCTTCATCAACCACACCCAAGGAATCCCAGACTTCTTCAAGCAATCCTTCCCAGAGGGATTCACCTGGGAGC
GGAAGTACATGCCTAGGGCATGGAAGTAGTTGGTGTGGGTTCCTTAGGGTCTGAAGAAGTTCGTTAGGAAGGGTCTCCCTAAGTGGACCCTCG
  94   96   98   100   102   104   106   108   110   112   114   116   118   120   122
S F M Y G S R T F I N H T Q G I P D F F K Q S F P E G F T W E
Translation
tagRFP
280 290 300 310 320 330 340 350 360 370
```

```
GTGTCACCACCTACGAGGACGGAGGAGTCCTCACCGCCACCCAAGACACCTCCCTCCAAGACGGATGCCTCATCTACAACGTCAAGATCCGTG
CACAGTGGTGGATGCTCCTGCCTCCTCAGGAGTGGCGGTGGGTTCTGTGGAGGGAGGTTCTGCCTACGGAGTAGATGTTGCAGTTCTAGGCAC
124   126   128   130   132   134   136   138   140   142   144   146   148   150   152   154
R V T T Y E D G G V L T A T Q D T S L Q D G C L I Y N V K I R
Translation
tagRFP
380 390 400 410 420 430 440 450 460
```

syb3140car2 recombined (891 bp) (from 466-891 bp)

GAGTCAACTTCCCATCCAACGGACCAGTCATGCAAAAGAAGACCCTCGGATGGGAGGCCAACACCGAGATGCTCTACCCAGCCGACGGAGGAC
CTCAGTTGAAGGGTAGGTTGCCTGGTCAGTACGTTTTCTTCTGGGAGCCTACCCTCCGGTTGTGGCTCTACGAGATGGGTCGGCTGCCTCCTG
156 158 160 162 164 166 168 170 172 174 176 178 180 182 184
G V N F P S N G P V M Q K K T L G W E A N T E M L Y P A D G G
Translation
tagRFP
470 480 490 500 510 520 530 540 550

TcGAGGGACGTACCGACATGGCCCTCAAGCTCGTCGGAGGAGGACACCTCATCTGCAACTTCAAGACCACCTACCGTTCCAAGAAGCCAGCCA
AgCTCCCTGCATGGCTGTACCGGGAGTTCGAGCAGCCTCCTCCTGTGGAGTAGACGTTGAAGTTCTGGTGGATGGCAAGGTTCTTCGGTCGGT
186 188 190 192 194 196 198 200 202 204 206 208 210 212 214 216
L E G R T D M A L K L V G G G H L I C N F K T T Y R S K K P A
Translation
tagRFP
560 570 580 590 600 610 620 630 640 650

AGAACCTCAAGATGCCAGGAGTCTACTACGTCGACCACCGTCTcGAGCGTATCAAGGAGGCCGACAAGGAGACcTACGTCGAGCAACACGAGG
TCTTGGAGTTCTACGGTCCTCAGATGATGCAGCTGGTGGCAGAgCTCGCATAGTTCCTCCGGCTGTTCCTCTGgATGCAGCTCGTTGTGCTCC
218 220 222 224 226 228 230 232 234 236 238 240 242 244 246
K N L K M P G V Y Y V D H R L E R I K E A D K E T Y V E Q H E
Translation
tagRFP
660 670 680 690 700 710 720 730 740

TCGCCGTCGCCCGTTACTGCGACCTCCCATCCAAGCTCGGACACAAGCTCAACGgTatggaTgaActGtaTaaGgtaagtttaaacatgattt
AGCGGCAGCGGGCAATGACGCTGGAGGGTAGGTTCGAGCCTGTGTTCGAGTTGCcAtacctActTgaCatAttCcattcaaatttgtactaaa
248 250 252 254 256 258 260 262 264 266 268 270 272 274 276 278
V A V A R Y C D L P S K L G H K L N G M D E L Y K V S L N M I
Translation
tagRFP
750 760 770 780 790 800 810 820 830

tactaacATAACTTCGTATAGCATACATTATACGAAGTTATAGttttttagcca
atgattgTATTGAAGCATATCGTATGTAATATGCTTCAATATCaaaaaatcggt
280 282
L L T *
Translation
840 850 860 870 880 890

**Figure S5. Sequence of sng-1(syb3140car2) cDNA after Cre/LoxP-mediated removal of GFP.**

(from 1-749 bp)

### sng-1 syb3140car2 genomic locus (2457 bp)

```
GCAACAACCACCATCAAACCCATAtACTCAaTCgGAAGGATATGGTTATGGAAGTTCCTATTCTCTAGAAAGTATAGGAACTTCATAAatttcaaattttaaatac
CGTTGTTGGTGGTAGTTTGGGTATaTGAGTtAGcCTTCCTATACCAATACCTTCAAGGATAAGAGATCTTTCATATCCTTGAAGTATTtaaagtttaaaatttatg
```

FRT | 3’ utr

10 20 30 40 50 60 70 80 90 100

```
tgaatatttgtttttttcctattatttatttattctctttgtgtttttttcttgctttctaaaaattaattcaatccaaatctaaacattttttttctctttc
acttataaacaaaaaaaggataataaataaataagagaaacacaaaaaaagaacgaaagatttttaattaagttaggtttagatttgtaaaaaaaagagaaag
```

3’ utr

110 120 130 140 150 160 170 180 190 200 210

```
cgtctcccaattcgtattccgctcctctcatctgaacacaatgtgcaagtttatttatcttctcgctttcatttcattaggacgtgggggaattggtggaagggggg
gcagagggttaagcataaggcgaggagagtagacttgtgttacacgttcaaataaatagaagagcgaaagtaaagtaatcctgcacccccttaaccaccttccccc
```

3’ utr

220 230 240 250 260 270 280 290 300 310 320

```
aaacacacaaaaggatgatggaaatgaaataaggacacacaatatgcaacaacattcaattcagaaatatggaggaaggtttaaaagaaaacataaaaatatataga
tttgtgtgttttcctactacctttactttattcctgtgtgttatacgttgttgtaagttaagtctttatacctccttccaaattttcttttgtatttttatatatct
```

3’ utr

330 340 350 360 370 380 390 400 410 420

```
ggaggaaggaaaactagtaaaaaataagcaaagaaattaggcgaacgatgagaattgtcctcgcttgGAAGTTCCTATTCTCTAGAAAGTATAGGAACTTCAGTCTC
cctccttccttttgatcatttttattcgtttctttaatccgcttgctactcttaacaggagcgaacCTTCAAGGATAAGAGATCTTTCATATCCTTGAAGTCAGAG
```

1 V, 2 S

3’ utr | FRT

430 440 450 460 470 480 490 500 510 520 530

```
CAAGGGAGAGGAGCTCATCAAGGAGAACATGCACATGAAGCTCTACATGGAGGGAACCGTCAACAACCACCACTTCAAGTGCACCTCCGAGGGAGAGGGAAAGCCAT
GTTCCCTCTCCTCGAGTAGTTCCTCTTGTACGTGTACTTCGAGATGTACCTCCCTTGGCAGTTGTTGGTGGTGAAGTTCACGTGGAGGCTCCCTCTCCCTTTCGGTA
```

3 K, 4 G, 5 E, 6 E, 7 L, 8 I, 9 K, 10 E, 11 N, 12 M, 13 H, 14 M, 15 K, 16 L, 17 Y, 18 M, 19 E, 20 G, 21 T, 22 V, 23 N, 24 N, 25 H, 26 H, 27 F, 28 K, 29 C, 30 T, 31 S, 32 E, 33 G, 34 E, 35 G, 36 K, 37 P

tagRFP

540 550 560 570 580 590 600 610 620 630 640

```
ACGAGGGAACCCAAACCATGCGTATCAAGGtaagtttaaacatatatatactaactaaccctgattatttaaattttcaggTCGTCGAGGGAGGACCACTCCCATTC
TGCTCCCTTGGGTTTGGTACGCATAGTTCCattcaaatttgtatatatatgattgattgggactaataaatttaaaagtccAGCAGCTCCCTCCTGGTGAGGGTAAG
```

38 Y, 39 E, 40 G, 41 T, 42 Q, 43 T, 44 M, 45 R, 46 I, 47 K

tagRFP

1 S, 2 S, 3 R, 4 E, 5 D, 6 H, 7 S, 8 H, 9 S

tagRFP

intron

650 660 670 680 690 700 710 720 730 740

sng-1 syb3140car2 genomic locus (2457 bp) (from 750-1391 bp)

```
GCCTTCGACATCCTCGCCACCTCCTTCATGTACGGATCCCGTACCTTCATCAACCACACCCAAGGAATCCCAGACTTCTTCAAGCAATCCTTCCCAGAGGGATTCAC
CGGAAGCTGTAGGAGCGGTGGAGGAAGTACATGCCTAGGGCATGGAAGTAGTTGGTGTGGGTTCCTTAGGGTCTGAAGAAGTTCGTTAGGAAGGGTCTCCCTAAGTG
10 11 12 13 14 15 16 17 18 19 20 21 22 23 24 25 26 27 28 29 30 31 32 33 34 35 36 37 38 39 40 41 42 43 44
P S T S S P P P S C T D P V P S S T T P K E S Q T S S S N P S Q R D S
tagRFP
750 760 770 780 790 800 810 820 830 840 850
```

```
CTGGGAGCGTGTCACCACCTACGAGGACGGAGGAGTCCTCACCGCCACCCAAGACACCTCCCTCCAAGACGGATGCCTCATCTACAACGTCAAGgtaagtttaaaca
GACCCTCGCACAGTGGTGGATGCTCCTGCCTCCTCAGGAGTGGCGGTGGGTTCTGTGGAGGGAGGTTCTGCCTACGGAGTAGATGTTGCAGTTCcattcaaatttgt
45 46 47 48 49 50 51 52 53 54 55 56 57 58 59 60 61 62 63 64 65 66 67 68 69 70 71 72 73 74 75
P G S V S P P T R T E E S S P P P K T P P S K T D A S S T T S
tagRFP
intron
860 870 880 890 900 910 920 930 940 950 960
```

```
gttcggtactaactaaccatacatatttaaattttcagATCCGTGGAGTCAACTTCCCATCCAACGGACCAGTCATGCAAAAGAAGACCCTCGGATGGGAGGCCAAC
caagccatgattgattggtatgtataaatttaaaagtcTAGGCACCTCAGTTGAAGGGTAGGTTGCCTGGTCAGTACGTTTTCTTCTGGGAGCCTACCCTCCGGTTG
1 2 3 4 5 6 7 8 9 10 11 12 13 14 15 16 17 18 19 20 21 22 23
I R G V N F P S N G P V M Q K K T L G W E A N
tagRFP
intron
970 980 990 1,000 1,010 1,020 1,030 1,040 1,050 1,060 1,070
```

```
ACCGAGATGCTCTACCCAGCCGACGGAGGACTcGAGGGACGTACCGACATGGCCCTCAAGCTCGTCGGAGGAGGACACCTCATCTGCAACTTCAAGACCACCTACCG
TGGCTCTACGAGATGGGTCGGCTGCCTCCTGAgCTCCCTGCATGGCTGTACCGGGAGTTCGAGCAGCCTCCTCCTGTGGAGTAGACGTTGAAGTTCTGGTGGATGGC
24 25 26 27 28 29 30 31 32 33 34 35 36 37 38 39 40 41 42 43 44 45 46 47 48 49 50 51 52 53 54 55 56 57 58 59
T E M L Y P A D G G L E G R T D M A L K L V G G G H L I C N F K T T Y R
tagRFP
1,080 1,090 1,100 1,110 1,120 1,130 1,140 1,150 1,160 1,170
```

```
TTCCAAGgtaagtttaaacatgattttactaactaactaatctgatttaaattttcagAAGCCAGCCAAGAACCTCAAGATGCCAGGAGTCTACTACGTCGACCACC
AAGGTTCcattcaaatttgtactaaaatgattgattgattagactaaatttaaaagtcTTCGGTCGGTTCTTGGAGTTCTACGGTCCTCAGATGATGCAGCTGGTGG
60 1 2 3 4 5 6 7 8 9 10 11 12 13 14 15 16
S K P A K N L K M P G V Y Y V D H
tagRFP
intron
1,180 1,190 1,200 1,210 1,220 1,230 1,240 1,250 1,260 1,270 1,280
```

```
GTCTcGAGCGTATCAAGGAGGCCGACAAGGAGACcTACGTCGAGCAACACGAGGTCGCCGTCGCCCGTTACTGCGACCTCCCATCCAAGCTCGGACACAAGCTCAAC
CAGAgCTCGCATAGTTCCTCCGGCTGTTCCTCTGgATGCAGCTCGTTGTGCTCCAGCGGCAGCGGGCAATGACGCTGGAGGGTAGGTTCGAGCCTGTGTTCGAGTTG
17 18 19 20 21 22 23 24 25 26 27 28 29 30 31 32 33 34 35 36 37 38 39 40 41 42 43 44 45 46 47 48 49 50 51 52
R L E R I K E A D K E T Y V E Q H E V A V A R Y C D L P S K L G H K L N
tagRFP
1,290 1,300 1,310 1,320 1,330 1,340 1,350 1,360 1,370 1,380 1,390
```

sng-1 syb3140car2 genomic locus (2457 bp) (from 1392-2033 bp)

```
GgTatggaTgaActGtaTaaGgtaagtttaaacatgattttactaacATAACTTCGTATAGCATACATTATACGAAGTTATtaactaatctgatttaaattttcagG
CcAtacctActTgaCatAttCcattcaaatttgtactaaaatgattgTATTGAAGCATATCGTATGTAATATGCTTCAATAattgattagactaaatttaaaagtcC
53 54 55 56 57 58 59
G  M  D  E  L  Y  K
tagRFP
intron   loxP   intron
1,400  1,410  1,420  1,430  1,440  1,450  1,460  1,470  1,480  1,490
```

```
GTTCTGGGAGTGGAAGCGGCTCTATGAGTAAAGGAGAAGAACTTTTCACTGGAGTTGTCCCAATTCTTGTTGAATTAGATGGTGATGTTAATGGGCACAAATTTTCT
CAAGACCCTCACCTTCGCCGAGATACTCATTTCCTCTTCTTGAAAAGTGACCTCAACAGGGTTAAGAACAACTTAATCTACCACTACAATTACCCGTGTTTAAAAGA
1 2 3 4 5 6 7 8 9 10 11 12 13 14 15 16 17 18 19 20 21 22 23 24 25 26 27 28
M S K G E E L F T G V V P I L V E L D G D V N G H K F S
GFP
GSGSGSGS linker
1,500  1,510  1,520  1,530  1,540  1,550  1,560  1,570  1,580  1,590  1,600
```

```
GTCAGTGGAGAGGGTGAAGGTGATGCAACATACGGAAAACTTACCCTTAAATTTATTTGCACTACTGGAAAACTACCTGTTCCATGGGTAAGTTTAAACATATATAT
CAGTCACCTCTCCCACTTCCACTACGTTGTATGCCTTTTGAATGGGAATTTAAATAAACGTGATGACCTTTTGATGGACAAGGTACCCATTCAAATTTGTATATATA
29 30 31 32 33 34 35 36 37 38 39 40 41 42 43 44 45 46 47 48 49 50 51 52 53 54 55 56 57
V S G E G E G D A T Y G K L T L K F I C T T G K L P V P W
GFP
intron
1,610  1,620  1,630  1,640  1,650  1,660  1,670  1,680  1,690  1,700  1,710
```

```
ACTAACTAACCCTGATTATTTAAATTTTCAGCCAACACTTGTCACTACTTTCTGTTATGGTGTTCAATGCTTCTCGAGATACCCAGATCATATGAAACGGCATGACT
TGATTGATTGGGACTAATAAATTTAAAAGTCGGTTGTGAACAGTGATGAAAGACAATACCACAAGTTACGAAGAGCTCTATGGGTCTAGTATACTTTGCCGTACTGA
1 2 3 4 5 6 7 8 9 10 11 12 13 14 15 16 17 18 19 20 21 22 23 24 25
P T L V T T F C Y G V Q C F S R Y P D H M K R H D
GFP
intron
1,720  1,730  1,740  1,750  1,760  1,770  1,780  1,790  1,800  1,810
```

```
TTTTCAAGAGTGCCATGCCCGAAGGTTATGTACAGGAAAGAACTATATTTTTCAAAGATGACGGGAACTACAAGACACGTAAGTTTAAACAGTTCGGTACTAACTAA
AAAAGTTCTCACGGTACGGGCTTCCAATACATGTCCTTTCTTGATATAAAAAGTTTCTACTGCCCTTGATGTTCTGTGCATTCAAATTTGTCAAGCCATGATTGATT
26 27 28 29 30 31 32 33 34 35 36 37 38 39 40 41 42 43 44 45 46 47 48 49 50 51
F F K S A M P E G Y V Q E R T I F F K D D G N Y K T
GFP
intron
1,820  1,830  1,840  1,850  1,860  1,870  1,880  1,890  1,900  1,910  1,920
```

```
CCATACATATTTAAATTTTCAGGTGCTGAAGTCAAGTTTGAAGGTGATACCCTTGTTAATAGAATCGAGTTAAAAGGTATTGATTTTAAAGAAGATGGAAACATTCT
GGTATGTATAAATTTAAAAGTCCACGACTTCAGTTCAAACTTCCACTATGGGAACAATTATCTTAGCTCAATTTTCCATAACTAAAATTTCTTCTACCTTTGTAAGA
1 2 3 4 5 6 7 8 9 10 11 12 13 14 15 16 17 18 19 20 21 22 23 24 25 26 27 28
A E V K F E G D T L V N R I E L K G I D F K E D G N I L
GFP
intron
1,930  1,940  1,950  1,960  1,970  1,980  1,990  2,000  2,010  2,020  2,030
```

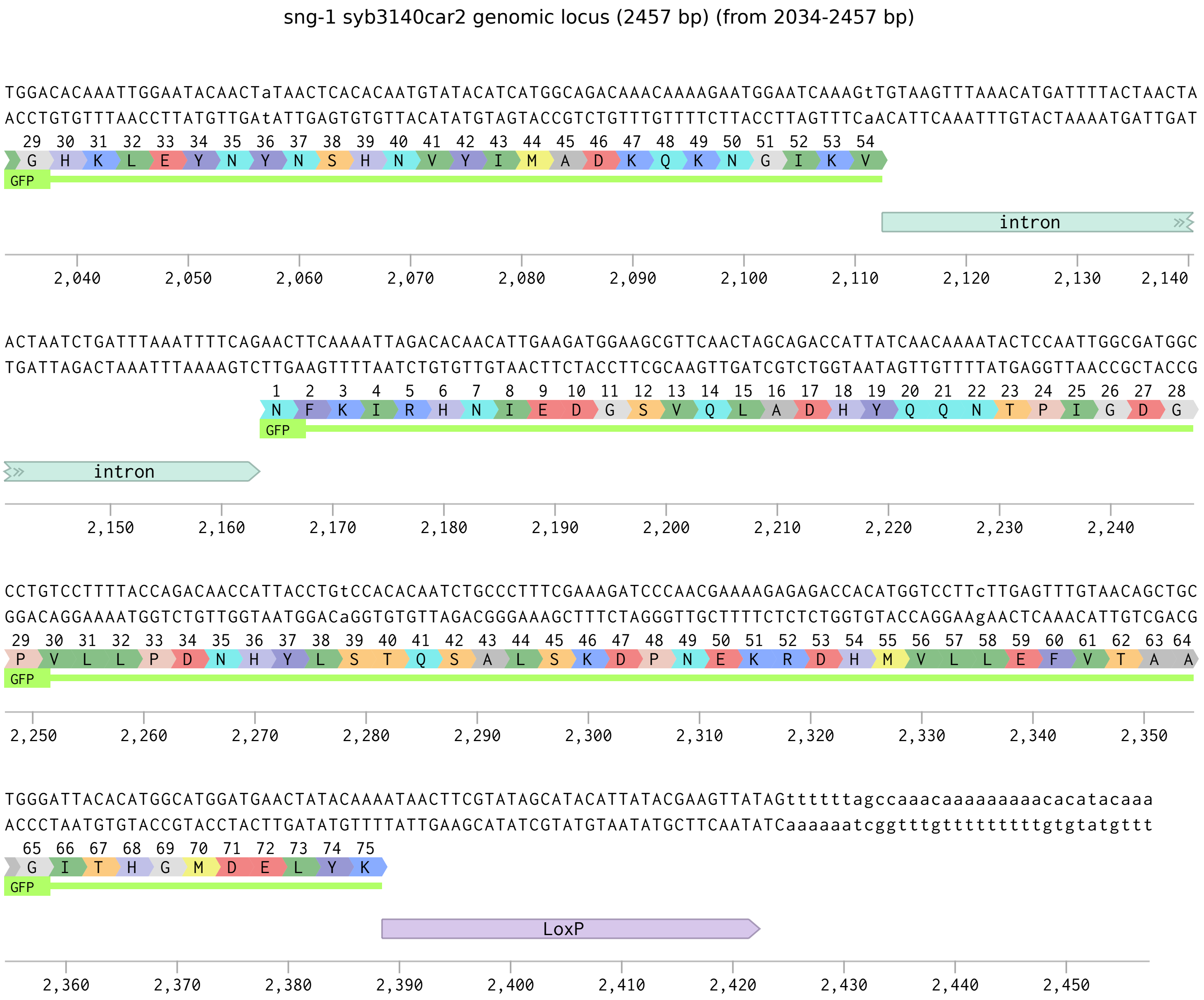


**Fig S6. Annotated genomic sequence of sng-1(syb3140car2[argo]).**
